# A qualitative and quantitative framework for the characterization of protein–RNA interactions by cross-linking mass spectrometry (XL-MS)

**DOI:** 10.64898/2026.10.15.756239

**Authors:** Izabela Smok, Aurélie Zbinden, Johannes Osterroth, Jianning Jiang, Frédéric H.-T. Allain, Jonathan Hall, Magdalini Polymenidou, Alexander Leitner

## Abstract

Protein–RNA interactions regulate nearly all levels of gene expression. Their characterization at the molecular basis is critical for understanding the function of protein–RNA complexes, both in cell homeostasis and pathology. Yet, molecular interactions underlying the formation of these complexes remain understudied. UV cross-linking mass spectrometry (XL-MS) is one of the leading approaches for *in vitro* identification of protein–RNA contact sites, and for providing structural information in the form of distance constraints crucial for the refinement of structural models. Our previously published CLIR-MS method enables qualitative characterization of protein–RNA interactions at residue-specific resolution but was not designed for highly accurate quantitative interpretation. Here, we present a XL-MS workflow based on irradiation with UV light that combines data-dependent acquisition (DDA) and parallel reaction monitoring (PRM) for simultaneous detection and characterization of protein–RNA interaction sites at residue-specific resolution, followed by label-free quantification of cross-linked products. We successfully apply our combined workflow to *in vitro* reconstituted protein–RNA complexes formed by the proteins TDP-43 and FOX1. We report on novel RNA interaction sites located in the oligomerization interface of the N-terminal domain (NTD) of TDP-43, and characterize quantitatively the process of RNA-induced monomerization of the NTD in an RNA titration experiment. We also investigate temperature-dependent differences in the efficiency of cross-link formation in the FOX1 complex. Our framework offers a unique and novel approach to comparative quantification of protein-RNA cross-links, overcoming the limitations imposed by conventional UV XL-MS methods.

## Introduction

Protein–RNA complexes are fundamental in the control and regulation of nearly all levels of gene expression.^1^ Understanding the molecular basis of protein–RNA interactions is critical for the comprehensive explanation of how proteins and RNAs assemble into complexes, bind their targets, and contribute to cellular homeostasis. The characterization of protein–RNA interactions is essential for the development of potential novel therapeutic targets.^2,3^ Protein–RNA interactions can be studied at large scale from two angles, either by identifying RNAs that are bound to a specific protein (protein-centric view) or by identifying proteins that are bound to a specific RNA (RNA-centric view). Cross-linking and mass spectrometry (XL-MS), as a protein-centric approach, is primarily used to identify RNA-binding proteins in complex samples, such as cell or tissue lysates, for example by employing organic-phase separation methods, like OOPS and XRNAX.^4,5^ XL-MS can be also used to identify nucleotide-interacting amino acids in *in vitro* reconstituted complexes at residue resolution.^6,7^ For studying protein–RNA interactions, XL-MS relies predominantly on ultraviolet (UV) light-induced formation of covalent bonds between amino acids and RNA nucleotides in close spatial proximity, followed by analysis of cross-linking products by mass spectrometry (MS). *In vitro* protein–RNA XL-MS has been also used for discovery purposes of novel RNA interaction sites in proteins^7–9^ or characterizing the photochemical selectivity of UV cross-links.^10^ XL-MS data can be used to generate distance restraints in integrative structural modeling,^11–13^ which is particularly useful for highly flexible structures, otherwise often difficult to study by conventional structural biology methods.^14–15^

Our previously published CLIR-MS (cross-linking of isotope labeled RNA coupled to mass spectrometry) method enables qualitative characterization of protein–RNA interactions at single-residue resolution *in vitro* but is not designed for quantitative interpretation. As a data-dependent acquisition (DDA) method, CLIR-MS is limited by the stochastic and abundance-biased process of precursor selection, which reduces measurement reproducibility and thus compromises quantitative accuracy.^16^ This is particularly limiting for peptide–RNA cross-links, often present in low abundance in a sample, even despite the enrichment procedure.^12^ In addition, transient or weak protein–RNA interactions, as well as their subtle changes may be difficult to characterize with a DDA-based workflow, especially in comparative analyses.

Here, we present a cross-linking workflow based on irradiation with UV light that combines data-dependent acquisition and parallel reaction monitoring (PRM) for simultaneous detection and characterization of protein–RNA interaction sites at residue-specific resolution, followed by label-free quantification of cross-linked products.

Our PRM-based label-free quantification relies on selective and sensitive detection of pre-defined targets (peptide–RNA cross-links, pre-identified using CLIR-MS) to provide lower detection limits, reduced background, and improved run-to-run reproducibility.^17–19^ By monitoring the abundance changes of selected cross-links, the method therefore allows the quantitative and reproducible description of the changes in protein–RNA interactions across various conditions. Normalization to an internal standard overcomes the variability of cross-link identification and site assignment across replicates,^12,20^ as well as allows direct comparison of samples across various experimental conditions, which is not achievable with DDA-based CLIR-MS.

Whereas CLIR-MS localizes a peptide–RNA cross-link through an approach analogous to that used for post-translational modifications, PRM-based analysis offers improved site assignment based on the extended identification of cross-link-specific fragment ions and RNA backbone ions. Moreover, because PRM provides insight into the chromatographic behavior of peptide–RNA cross-links, it may offer a means to distinguish isobaric products (positional isomers, neutral loss products) arising from UV light irradiation and sample processing.

We applied our combined DDA-PRM workflow to a model system, human TAR DNA-binding protein 43 (TDP-43) in a complex with specifically recognized^21^ (GU)_6_ RNA, and human RNA binding protein fox-1 homolog 1 (FOX1) in a complex with specifically recognized^22^ FOX1 binding element (FBE) RNA. With this method, we improve the confidence of cross-link site localization on both peptide and RNA by predicting the fragment ions of peptide–RNA adducts in positive mode. We identify additional novel RNA interaction sites localized in the oligomerization interface of the TDP-43 NTD protein and demonstrate that RNA interactions stabilize TDP-43 NTD monomers by quantifying the changes of cross-link abundances upon adding increasing amounts of RNA. We also investigate how sample temperature influences the efficiency of UV cross-link formation in the FOX1–FBE complex. The presented approach provides a novel strategy for comparison of protein-RNA interactions across biophysical and structural conditions, and addresses and overcomes the limitations of conventional UV XL-MS methods.

## Results

### Chymotrypsin-based digestion reveals additional RNA interaction sites in the oligomerization interface of TDP-43 NTD

Human TDP-43 (Fig. 1A) is a ubiquitously expressed, primarily nuclear RNA-binding protein (RBP), involved in mRNA translation and stability, mitochondrial and synaptic homeostasis and stress responses.^23–25^ Upon physiological conditions, TDP-43 predominantly localizes to the nucleus, while its increased cytoplasmic accumulation and aggregate formation in neurons have been correlated with neurodegenerative diseases, such as amyotrophic lateral sclerosis (ALS) and frontotemporal dementia (FTD).^26,27^

**Figure 1.**
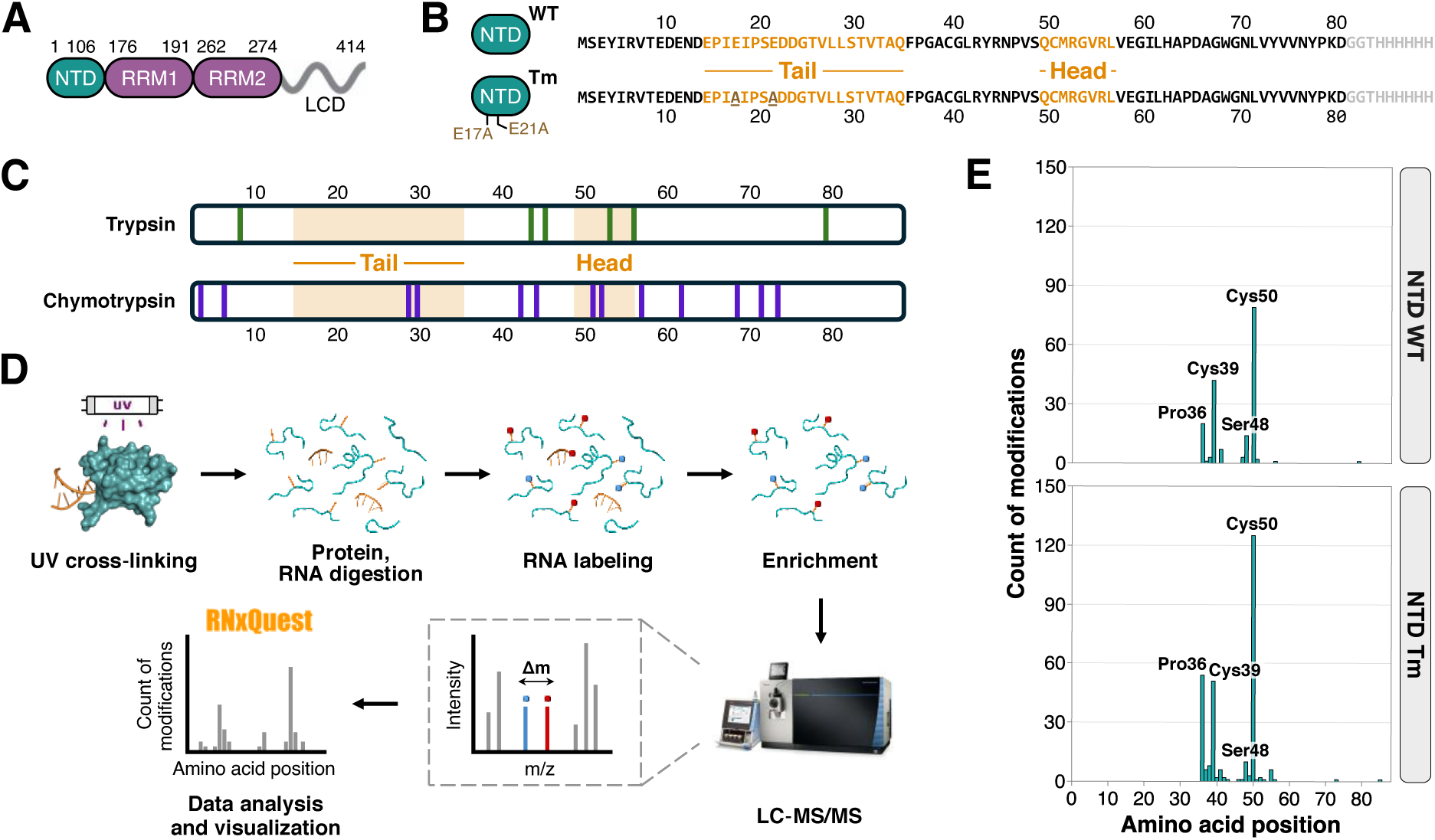
Chymotrypsin-based digestion reveals additional RNA interaction sites in the oligomerization interface of TDP-43 NTD. **(A)** Schematic representation of human TDP-43 domain architecture. NTD: N-terminal domain, RRM: RNA-recognition motif, LCD: low-complexity domain. **(B)** Sequence overview of TDP-43 NTD variants used in the experiment. Residues representing the oligomerization interface are highlighted in orange, the C-terminal 6xHis-tag is highlighted in grey. Mutated residues in the oligomerization-deficient variant, NTD Tm, are highlighted in brown. **(C)** Comparison of sequence coverage of TDP-43 NTD. Trypsin (KR/P) and chymotrypsin cleavage sites (FWYML/P) are denoted as vertical bars. The residues representing the oligomerization interface are highlighted in orange. **(D)** Schematic representation of the CLIR-MS workflow. *In vitro* reconstituted protein–RNA complex is irradiated with UV light at 254 nm and subsequently digested with RNases and a protease. Cross-linked RNA is subject to enzymatic ^18^O labeling. Enriched peptide–RNA cross-links are separated by LC and ionized in the mass spectrometer, and their mass-to-charge (*m/z*) is determined. In precursor mass scans (MS1), only the peptides that were cross-linked to the labeled RNA segment appear as “doublets” with a distinct mass shift (Δm), introduced by the ^18^O labeling step. Abundant peptide–RNA cross-links are fragmented (MS2) to determine the position and composition of the covalently linked RNA adduct. Data is processed and visualized with xQuest and RNxQuest software. **(E)** Total number of detected cross-links (peptide-spectrum-matches, PSMs) identified for TDP-43 NTD WT–(GU)6 complex (top) and TDP-43 NTD Tm–(GU)6 complex (bottom), generated with the CLIR-MS workflow. Identified cross-links are mapped onto the amino acid positions of the respective protein variant.

TDP-43 is composed of two RNA recognition motifs (RRMs), responsible for specific recognition of nucleic acids enriched in TG-/UG repeats,^25,28^ a C-terminal intrinsically disordered domain (low-complexity domain, LCD), mediating protein-protein interactions and liquid-liquid phase separation (LLPS), and an N-terminal domain (NTD), facilitating protein oligomerization. TDP-43 oligomerizes in a concentration-dependent manner, upon head-to-tail interactions between the negatively charged “tail” region (E14–Q34) and the positively charged “head” region (Q49–L56) of two monomers^29^ (Fig. 1B). Higher oligomeric species are stabilized further by intermolecular disulfide bridges.^30^ Physiological TDP-43 oligomerization has been shown as crucial for numerous cellular functions, such as subcellular protein localization, LLPS and alternative splicing, highlighting its importance in pathology.^8,29,31–33^ Recently, we have shown that TDP-43 NTD interacts with (GU)_6_ RNA, independently of the RRMs of the full-length protein.^8^ Using CLIR-MS workflow, we identified a novel RNA cross-linking site at Cys50, located at the oligomerization interface of the protein. This finding suggested a functional interplay between RNA interaction and protein oligomerization. To better characterize the dynamic relationship between these two processes, we sought to first improve the sequence coverage of the TDP-43 NTD to uncover additional RNA interaction sites.

Comprehensive characterization of protein–RNA interaction sites in XL-MS relies heavily on efficient digestion of protein and RNA. Trypsin remains the protease of choice in conventional bottom-up proteomics workflows, owing to its well-defined specificity and high activity. It cleaves C-terminal to Lys and Arg residues, resulting in peptides that usually contain at least two positive charges at both termini, granting efficient ionization and fragmentation in the mass spectrometer.^34^ However, MS for structural biology often incorporates alternative proteases, for improving the detection of cross-links.^35–37^

In our earlier study,^8^ trypsin was successfully used to investigate RNA interaction sites of the NTD of TDP-43 protein, in complex with (GU)_6_ RNA. However, trypsin treatment resulted in suboptimal cleavage, due to general lack of Lys and Arg residues in the NTD (Fig. 1C). Therefore, to build on our previous trypsin-based findings, we chose chymotrypsin as an alternative protease to broaden the sequence coverage of TDP-43 NTD. Chymotrypsin primarily targets carboxyl sides of Phe, Tyr, Trp, Met and Leu,^38^ offering improved sequence coverage of the oligomerization interface, and thus potentially revealing more potential protein– RNA interaction sites.

Similarly to our previous study, the DDA-based CLIR-MS workflow (Fig. 1D) was first applied to two TDP-43 NTD variants, wild type (NTD WT) and oligomerization-deficient tail mutant (NTD Tm, E17A/E21A; Fig. 1B), both independently in a complex with (GU)_6_ RNA. Samples were treated with chymotrypsin instead of trypsin, while other protocol steps remained unchanged.

Our results uncover additional protein–RNA interaction sites, not detected in the previous, trypsin digestion-based experiment, with Pro36 and Cys39 being the interaction hotspots (Fig. 1E). Both novel sites map to the peptide 29-STVTAQFPGACGL-41, identified both in its fully cleaved form and with extensions on the N-and C-termini due to miscleavages. These interaction sites have not been observed with trypsin digestion, most likely due to the length of the corresponding tryptic peptide (over 30 residues). Both Pro36 and Cys39 are detected independent of the protein variant used, suggesting that oligomeric state of TDP-43 NTD does not qualitatively influence RNA interactions. Cys39 of TDP-43 NTD WT has been shown previously to undergo substantial chemical shift perturbations (CSPs) upon increasing protein concentration, therefore suggesting its role in protein oligomerization.^29^

Importantly, our chymotrypsin-based experiment also detects a second RNA interaction cluster, with Cys50 as a dominant site and Ser48 as a secondary site. Both map to the peptide 44-RNPVSQCM-51 and its variants with miscleavages, in the head region of protein’s oligomerization interface. This strengthens our previous finding^8^ and shows the detection consistency of the functional interplay between TDP-43 oligomerization and RNA interactions via a shared interface.

All protein–RNA cross-links were detected with GU as the attached RNA composition, independent of the protein variant complexed to (GU)_6_. This also confirms our previous findings and additionally indicates no difference in preferred RNA composition recognized and bound by two separate cross-linking clusters, Pro36/Cys39 and Ser48/Cys50.

### TDP-43 NTD interacts with RNA via Cys39 and Cys50 residues, independent of protein oligomeric state

The established CLIR-MS method supports cross-link site localization, up to a single peptide residue and a single nucleotide. Yet, often the desired resolution is not achievable, mainly owing to the previously mentioned abundance bias and low spectral quality. Additionally, RNA-level single nucleotide resolution at the cross-link level depends on the cleavage efficiency of RNases used – any product RNA left off as part of a cross-link longer than one nucleotide is reported as an RNA composition (but not the sequence), without specifying the exact nucleotide involved in cross-link formation.

TDP-43 NTD cross-links initially detected by DDA-based CLIR-MS workflow (Fig. 1E) are also subject to this limitation. For example, Pro36 and Cys39 are both assigned as dominant RNA interaction sites, alongside several less pronounced adjacent residues — all mapping to the same 29-STVTAQFPGACGL-41 peptide. Targeted acquisition is a method of choice for distinguishing between two possible scenarios: cross-linking site misassignment of residues belonging to the same peptide, and coexistence of multiple species, each cross-linking to RNA through a distinct amino acid residue of the same peptide.

Therefore, we targeted selected cross-links (chymotryptic peptide–RNA adducts detected by CLIR-MS), reporting on two dominant interaction clusters observed, Pro36/Cys39 and Ser48/Cys50 with our PRM-based approach (Fig. 2A). To pinpoint the exact cross-linked residue of a peptide, we performed spectrum-centric matching of the acquired MS2 spectra of each targeted cross-link against an *in silico* transition list of predicted transitions. To ensure unbiased site localization, we considered cross-link position at every candidate residue across all peptide–RNA adducts targeted in the PRM experiment. Full series of theoretical b-and y-ions were generated, including unmodified (non-cross-linked to intact GU RNA) and modified (cross-linked to intact GU RNA) peptide backbone fragments.

**Figure 2.**
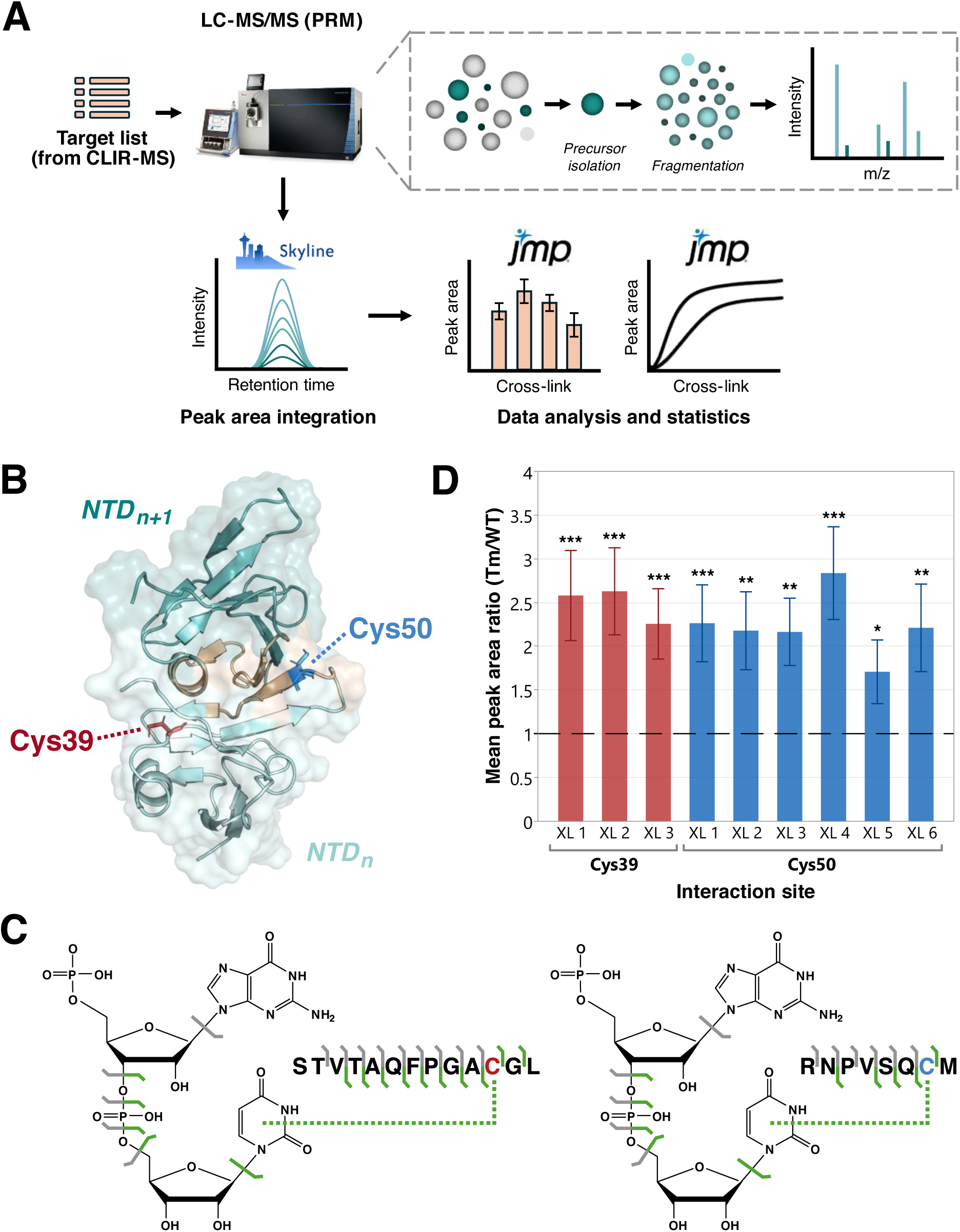
GU RNA interactions are enhanced in the monomeric state of TDP-43 NTD. **(A)** Schematic representation of the PRM workflow. Target peptide–RNA cross-links (pre-identified using CLIR-MS) are selectively isolated and fragmented to yield full fragmentation spectra. The chromatographic traces of co-eluting fragment ions (MS2) are extracted using Skyline software for peak area integration. Normalization to an internal standard and statistical analysis is performed with JMP Student software. **(B)** TDP-43 NTD dimer (PDB 6B1G)^33^ structure with mapping of detected interaction sites. Oligomerization interface is highlighted in yellow. **(C)** Annotation of observed fragment ions on the proposed structure of peptide–RNA cross-links carrying a single phosphate group at the 5′ end. Dominant cross-link structure reporting on the Cys39 site is shown on the left, the dominant cross-link for the Cys50 site is shown on the right. Green bars mark fragment ions (MS2) reporting on cross-link between peptide and RNA, grey bars – non-cross-linked fragment ions. **(D)** Fold change observed between the oligomeric NTD WT and oligomerization-deficient NTD Tm, shown as a ratio of individually integrated peak areas per cross-link type (cross-links specified in the Suppl. Tab. 1). The significance of abundance changes was assessed using a paired *t*-test, with p-values adjusted for multiple comparisons by the Benjamini–Hochberg procedure (* for q < 0.05, ** for q < 0.01, *** for q < 0.001). Error bars represent the standard error of the mean (SEM).

We were thus able to pinpoint Cys39 and Cys50 as strongly dominant RNA cross-linking sites (Fig. 2B). All other residues (including Pro36 and Ser48) are either very low abundant products or misassigned by the software due to poor spectral quality or co-fragmentation, whereas PRM improves localization confidence through repeated fragmentation of targeted cross-links. Our observation is consistent among peptides with missed cleavages and different 3′/5′ phosphorylation states of the cross-linked GU dinucleotide. The cysteine-based interaction was found to be independent of oligomeric state of protein.

While our previous MS experiments already revealed Cys50 as a novel RNA interaction site of TDP-43, Cys39 has not been reported before in this context.^8^ However, both have been shown in the literature to orchestrate tetramerization of TDP-43 by formation of intermolecular disulfide bridges,^30^ disrupted upon double mutation to serine residues (C39S/C50S).^33^

### Detailed analysis of PRM data allows detection of the cross-linked **nucleobase and the 5′ to 3′ order of nucleotides**

Based on the same approach used for pinpointing the cross-linked peptide residue, we sought to elucidate the specific cross-linked nucleotide within the detected GU RNA composition. We generated an alternative *in silico* transition list, fixing the cysteines as the cross-linked residues and predicted full series of RNA transitions (3′ end-retaining w-, x-, y-, z-ions; 5′ end-retaining a-, b-, c-, d-ions; nucleobase, ribose), according to the standard RNA fragmentation nomenclature.^39^ RNA transitions included unmodified (non-cross-linked to peptide backbone) fragments and modified (cross-linked to peptide backbone) fragments. Since the detected RNA composition was GU, both G and U nucleotides were considered as potential cross-linked moieties.

We successfully identified uracil as the directly interacting RNA moiety, by observing transitions corresponding to cross-linked U base to peptide backbone fragment ions (Fig. 2C). We also detected a singly charged G base fragment ion (152 m/z), predominantly as the most intense signal observed in the MS2 spectra. No U base fragment ion was observed (113 m/z). Our findings are consistent among all PRM-targeted cross-links. Preferential cross-link formation to U base is shared by both Cys39 and Cys50, independent of oligomeric state of the protein.

Structural mapping of detected fragment ions (Fig. 2C) reveals the predominantly observed structures of peptide–RNA cross-links. Both Cys39 and Cys50 cross-link to the 3′-terminal U base, suggesting 5′-GU-3′ as the dominant RNA sequence bound. It remains to be determined whether 5′-GU-3′ arises from the RNA sequence recognition by TDP-43 NTD, or from structural geometry, favoring cross-link formation. The observed 5′ phosphorylation matches the expected outcome of the ^18^O labeling procedure.^6^

### GU RNA interactions are enhanced in the monomeric state of TDP-43 NTD

While both complexes show no difference in preferred interaction sites and RNA compositions revealed by CLIR-MS (Fig. 1E), oligomerization-deficient NTD Tm exhibits a great increase in detected cross-links compared to oligomerizing NTD WT, particularly pronounced at the Cys50 cross-linking site. The effect is, to much lesser extent, also observed at the Cys39 site. This results hint that the two interaction sites may differ in their RNA interaction responsiveness, governed by the oligomeric state of the protein.

The DDA-based CLIR-MS workflow is, however, not sufficient to provide robust quantification of observed effects. Semi-quantitative profiling performed by the RNxQuest software relies on spectral counting, in which the number of assigned fragment ion (MS2) spectra serves only as a proxy for abundance. Alternatively, DDA-based quantification of peptide–RNA cross-links can be performed at the precursor (MS1) level, which involves the integration of precursor peak areas and signal normalization between runs. This approach, however, remains susceptible to interferences, such as the co-elution of positional isomers or of neutral loss products of the same cross-link. Both approaches are limited by the stochastic nature and limited sensitivity of DDA acquisition, in which precursors are sampled inconsistently across runs. Therefore, we applied our PRM-based approach to perform label-free quantification at the MS2 level. This allows to directly compare the extent of RNA interactions at two individual cross-linking sites, observed in two protein variants. We relied on the same dataset used for assessing site localization on both peptide-and RNA-level.

Oligomerization-deficient NTD Tm shows significant, over 2-fold increase in cross-link formation with GU RNA composition, compared to oligomeric NTD WT (Fig. 2D). This is consistent with our previous finding,^8^ emphasizing the increased RNA binding upon protein monomerization: the oligomerization interface, inaccessible in the assembled state, becomes surface-exposed in the monomer, allowing RNA to interact. Over 2-fold increase implies that dimers and trimers are the dominant oligomeric species formed by the wild-type protein variant, assuming a discrete monomerization process. Our initial data showed over 3-fold increase, most likely related to even higher oligomeric species present back then in the analyzed samples.

The observed trend is consistent across both interaction sites, implying that both sites show a comparable degree of response to RNA interactions upon protein monomerization, independent of their solvent accessibility – despite Cys39 being more surface-exposed than Cys50 (20% and 7% relative solvent accessibility, respectively, based on PDB 6B1G).^33^

Owing to its deterministic acquisition, our targeted approach therefore provides a more reliable and reproducible measure of the differences in RNA interactions between the two protein variants than the abundance-biased DDA-based CLIR-MS approach. Remarkably, the fold changes are lower than initially suggested by CLIR-MS, particularly at the Cys50 cross-linking site.

### Specific RNA interactions trigger the dissociation of TDP-43 NTD oligomers

The enhanced interaction of the oligomerization-deficient NTD Tm with (GU)_6_ RNA suggested that both protein oligomerization and RNA binding compete for the same TDP-43 interface and rely on similar electrostatic interactions.^8^ We therefore hypothesized that RNA binding may induce TDP-43 monomerization.

To test this hypothesis, we performed a UV cross-linking-based experiment in which (GU)_6_ RNA was titrated into oligomeric NTD WT of a fixed concentration, across increasing RNA-to-protein ratios, and used Native PAGE as an initial readout (Fig. 3A). We observed a gradual, RNA-dependent shift from the oligomeric to the monomeric and RNA-complex forming species. This transition noticeably emerges above a 0.75 RNA-to-protein ratio, suggesting that near-stoichiometric interaction is sufficient for simultaneous oligomer dissociation and protein–RNA complex formation. Bands corresponding to stable monomeric species gradually intensify above the transition point, suggesting that the threshold concentration required for oligomer formation was not reached under these conditions.

**Figure 3.**
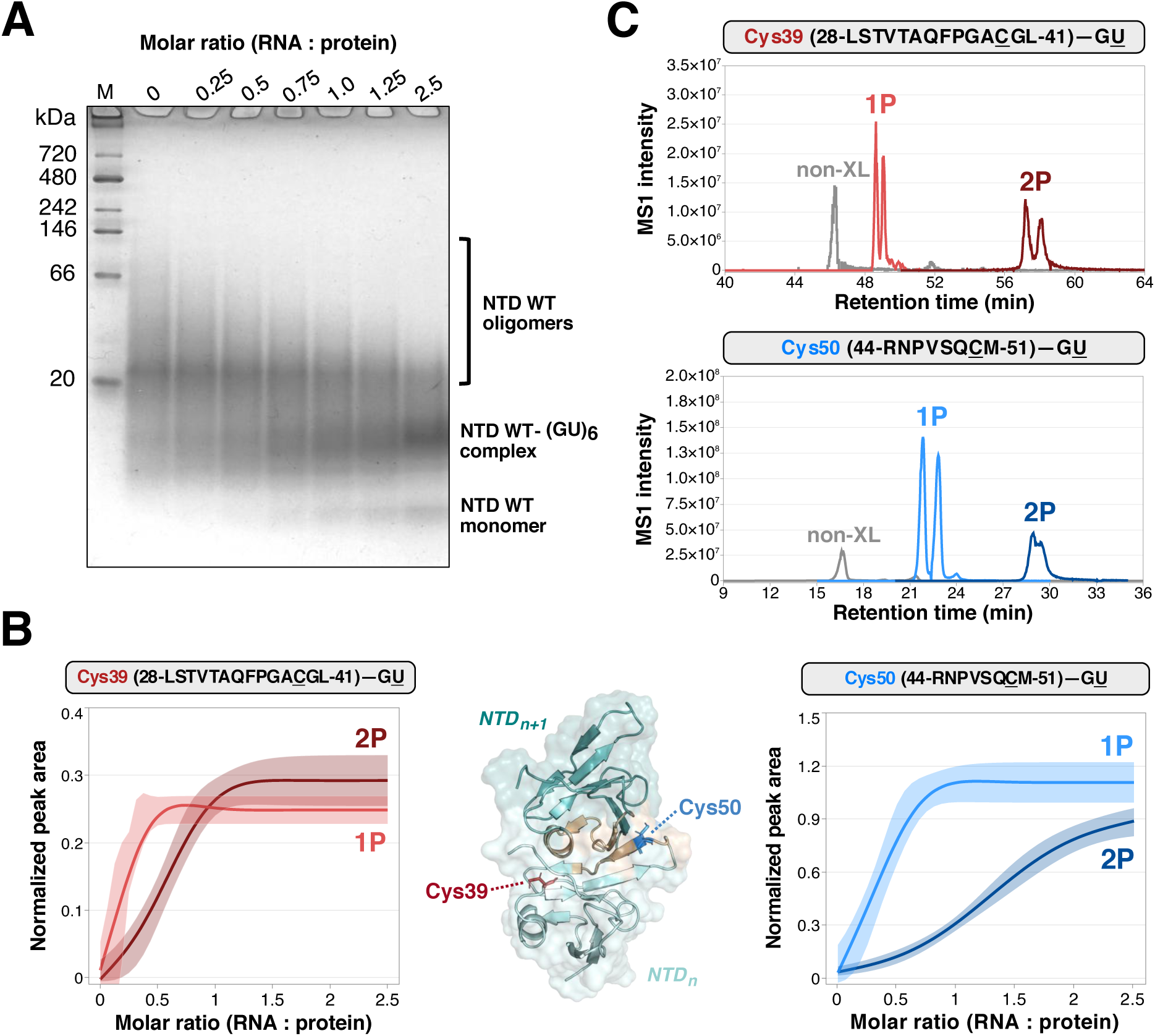
(GU)6 RNA titration of TDP-43 NTD WT shifts the equilibrium to the monomeric form. **(A)** Native PAGE separation of the UV cross-linked NTD WT–(GU)6 across increasing RNA-to-protein ratios. M denotes the molecular weight standard. **(B)** Titration curves obtained for the cross-links reporting on the Cys39 (left) and on the Cys50 interaction site (right), present as pairs of the phosphate-lacking (1P) and phosphate-retaining (2P) RNA forms. Peptide and RNA residues involved in cross-link formation are underlined. GU denotes the RNA composition, rather than the sequence. Normalized peak area data points across RNA–protein molar ratios were fitted to a three-parameter equation. The 95% confidence band around each fit represents mean predicted response. **(C)** Retention time comparison of non-cross-linked peptide (grey) and the cross-linked forms of the same peptide with RNA lacking one phosphate group (1P, light red or light blue) or retaining it (2P, dark red or dark blue). Peptide and RNA residues involved in cross-link formation are underlined. GU denotes the RNA composition, rather than the sequence. Data shown on the example of one of the replicates of RNA-to-protein ratio of 1. The observed retention pattern is consistent with the other measured RNA-to-protein ratios.

To quantitatively characterize this process at the residue level, we applied our PRM-based workflow, following the same RNA-to-protein ratios used for the Native PAGE. We reasoned that a progressive increase in the monitored peptide–RNA cross-link signals arises from increased oligomerization interface accessibility, as a result of protein monomerization.

All targeted peptide–RNA cross-links, regardless of interaction site, showed a significant increase in abundance of peptide–RNA cross-links upon (GU)_6_ titration, consistent with RNA-induced stabilization of protein monomers (Suppl. Fig. 1). Protein-protein interactions in the oligomerization domain may be outcompeted by multivalent interactions driven by RNA, including electrostatic interactions, hydrogen bonds and π-stacking. The individual transition points observed suggest weak affinity of NTD monomer self-association, in agreement with previous reports.^29,33.40,41^ 12-mer RNA sequence proves sufficient for robust oligomer disruption.

Importantly, the rate of the observed effect differs between the two RNA-interacting sites: Cys39 consistently displayed lower inflection points than those at Cys50, among all targeted cross-links (Fig. 3B). This suggests that NTD WT engages RNA interaction more effectively at the more surface-exposed Cys39, whereas the buried Cys50 residue requires higher molar excess of (GU)_6_ to promote protein monomerization and subsequent RNA interaction.

Interestingly, we also observed consistent differences in titration behavior of two RNA forms of the same peptide that differ by a single phosphate group: the phosphate-retaining (5′ and 3′ end of GU phosphorylated, 2P) and the phosphate-lacking form (either 5′ or 3′ end of GU phosphorylated, 1P). At both interaction sites, the phosphate-lacking form consistently yielded 3-fold lower fitted inflection points than the phosphate-retaining form. Both forms are generated downstream of UV cross-linking step, upon post-digest ^18^O labeling, which is catalyzed by T4-PNK enzyme.^6^ T4-PNK exhibits 5′-kinase activity, and a secondary 3′-phosphatase activity, making the 3′ phosphate-lacking form the intended product of RNA labeling, and the phosphate-retaining the result of incomplete 3′ phosphate removal. Since the enzymatic reaction is performed after protease and RNase digestions, the two forms are chemically identical at the moment of cross-link formation and differ only in the completeness of the 3′ dephosphorylation step.

The phosphate-retaining and phosphate-lacking forms are consistently well resolved chromatographically. The phosphate-retaining form elutes substantially later than its phosphate-lacking counterpart (up to 9 minutes offset at Cys39 site and up to 7 minutes at Cys50) (Fig. 3C). Increased retention of phosphorylated peptides in the presence of 0.1% formic acid has been reported previously,^42,43^ where the observed shift was even more pronounced for doubly-phosphorylated peptides. The effect can be attributed to ion pairing of the phosphate group with formate counterions present in the mobile phase; phosphopeptides therefore behave as more hydrophobic and are retained longer than the unmodified equivalent. This mechanism alone, however, does not account for the magnitude of the shift observed by us.

Cross-linked peptides show consistent chromatographic peak splitting, independent of the number of phosphate groups present (Fig. 3C). Peaks predominantly split into two partially resolved species that are uniform at the MS2 level, in intensity and peak area of detected fragment ions. One possible explanation of this observation is cis/trans isomerization of the Xaa–Pro peptide bond, enhanced by the bound RNA. We also observe a differential behavior of a miscleaved 42-RYRNPVSQCM-51 peptide that shows minor peak splitting in the non-cross-linked form and pronounced in the cross-linked form (up to 4 partially resolved species, Suppl. Fig. 2). This effect is also phosphate-dependent: the phosphate-retaining form yields three peaks and the phosphate-lacking form yields four, with the last two peaks in each group showing hindered peptide backbone fragmentation regardless of phosphate number. We attribute the phosphate dependence to additional interactions that lock a specific proline conformer, most likely through the previously proposed arginine forks between the N-terminal Arg and terminal phosphate groups. Other explanations of chromatographic peak splitting, such as positional isomers arising from RNA cross-linking to different positions in the amino acid side chain, or from peptide cross-linking to different atomic positions of the nucleobase, cannot be ruled out.

### Specific RNA recognition by FOX1-RRM protein is temperature-independent

We also applied our combined DDA-PRM workflow to another model system, the RRM domain of human FOX1 protein (Fig. 4A) in a complex with its specifically recognized FBE (5′-UGCAUGU-3′) RNA. Human FOX1 is an RNA splicing factor that regulates gene expression in heart, muscle and neuronal tissues during early development. FOX1 contains a single RRM domain that specifically recognizes and binds the (U)GCAUG sequences.^22^ In the nucleus, FOX1 regulates alternative splicing by recognizing the introns flanking alternative exons, whereas in the cytoplasm it influences gene expression by binding the 3′-untranslated regions (3′UTRs) of its target genes. Mutations in the FOX1 gene have been associated with autism-spectrum disorders (ASDs), intellectual disability and epilepsy syndromes.^44,45^

**Figure 4.**
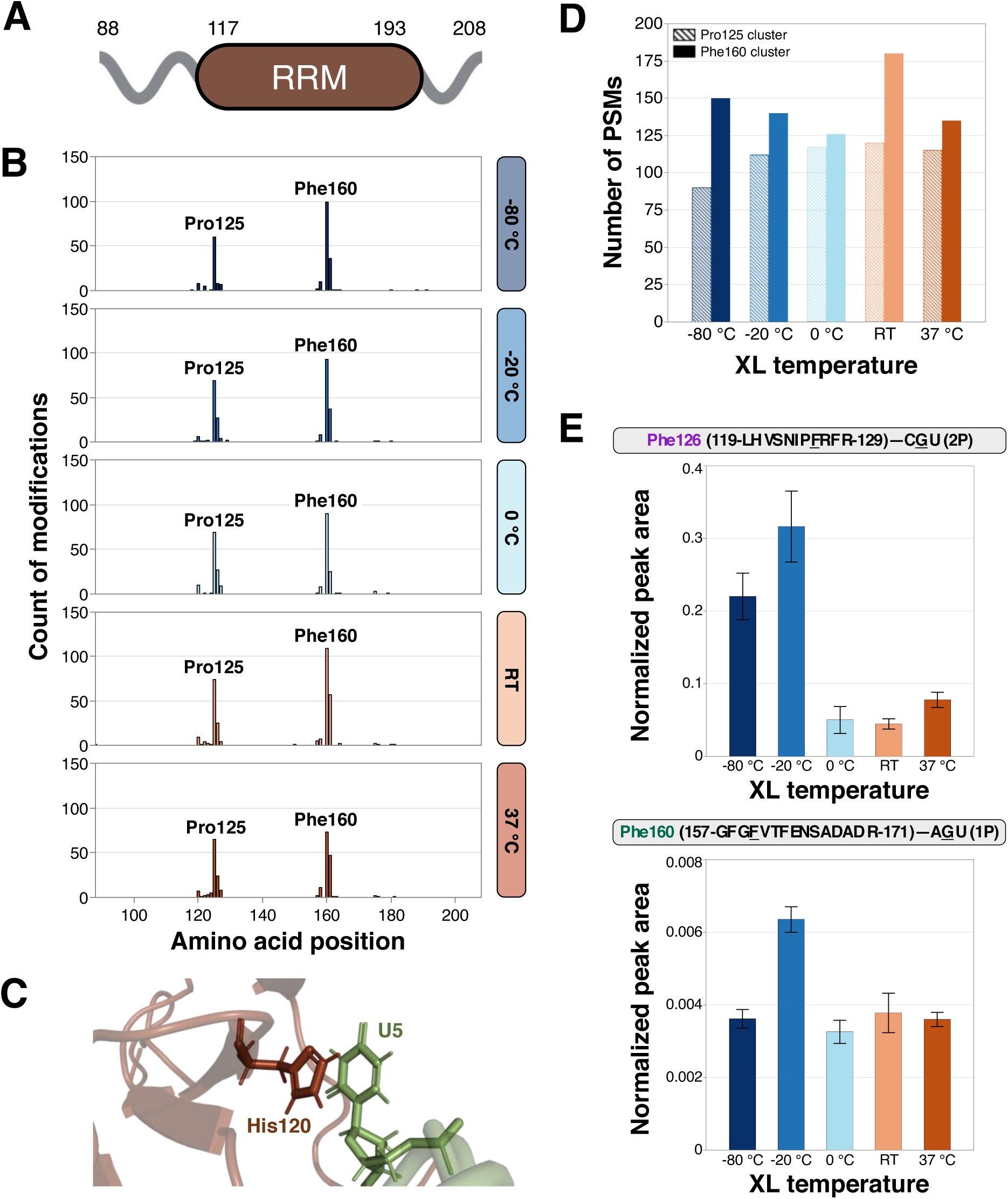
UV cross-linking of FOX1-RRM–FBE complex in the frozen state indicates improved reaction efficiency. **(A)** Schematic representation of domain architecture of the human FOX1 variant (FOX1-RRM). RRM: RNA-recognition motif. **(B)** Total number of detected cross-links identified for FOX1-RRM–FBE complex across different sample temperatures maintained during UV cross-linking procedure, generated with the CLIR-MS workflow. Identified cross-links are mapped onto the amino acid positions of FOX1-RRM. **(C)** Zoomed-in structure of FOX1-RRM–FBE complex (PDB 2ERR)^22^ showing stacking of His120 and U5 residues. **(D)** Comparison of peptide-spectrum-matches (PSMs) derived from the CLIR-MS workflow, reflecting the number of detected cross-links in two main interacting clusters observed, across conditions. Values represent data pooled across two technical replicate injections. **(E)** Comparative quantification of two selected cross-links showing the difference in peptide-RNA cross-link abundance across conditions. Peptide and RNA residues involved in cross-link formation are underlined. CGU and AGU denote the RNA composition, rather than the sequence. Error bars represent the standard error of the mean (SEM); n = 4.

Previously, we performed a DDA-based CLIR-MS workflow on the FOX1-RRM– FBE complex to evaluate the effect of UV cross-linking temperature on the number of identified cross-links. While the detected protein interaction sites remained unchanged across conditions, the number of cross-links at each interaction site showed temperature-dependent trends.^12^ We sought to investigate the observed differences quantitatively using the DDA-PRM workflow.

Solutions of the *in vitro* reconstituted FOX1-RRM–FBE complex were irradiated with the same total energy at 254 nm, but the UV cross-linking procedure was performed on samples at varying temperatures, including a sample heated to 37 °C, at room temperature (RT), cooled down on ice (approximately 0 °C), and frozen at-20 °C and-80 °C. All conditions show consistent identification of the two dominant RNA interaction sites, Pro125 and Phe160 (Fig. 4B). This suggests that RNA recognition is specific and governed by the conformationally stable interface of FOX1-RRM.

Interestingly, this specific recognition is not influenced by the potential acidic pH shift of sodium phosphate buffer during slow freezing.^46^ This is particularly important owing to the presence of His120 in the first interaction cluster. At low pH, histidine becomes positively charged, thus disrupting any existing hydrogen bonds and favoring electrostatic interactions. His120 has been observed to stack with the U5 nucleotide of FBE RNA (PDB 2ERR,^22^ Fig. 4C). His120 protonation may therefore convert the neutral π-stacking into a cation-π interaction, preserving the molecular contact. Alternatively, even if His120–FBE is disrupted, cross-links may still form through interactions driven by adjacent protein residues, such as Phe126 protein variants.^10^ We relied on the same dataset used for assessing site localization on both peptide-and RNA-level.

By comparing the PSMs derived from the CLIR-MS experiment, reflecting the number of detected cross-links across temperatures during UV irradiation, we observed minor differences between the two dominant interaction sites (Fig. 4D). In the first interaction cluster (Pro125), we observed a comparable number of identifications across all conditions, except for the-80 °C sample temperature, which yielded approximately 20% fewer PSMs. In the second interaction cluster (Phe160), we detected the highest number of identifications in the RT condition, while the remaining conditions yielded comparable numbers, approximately 20% fewer PSMs than at RT. The highest number of detected cross-links in both interactions clusters is observed in the RT condition, whereas the other conditions yielded comparable outcomes. The lowest applied temperature (-80 °C) resulted in the highest ratio of cross-links detected in the Phe160 cluster to those in the Pro125 cluster. This observation may further indicate an effect of His120 protonation on cross-links formation. Conventional UV irradiation temperature (0 °C) yielded a comparable number of cross-links relative to the other conditions, equally distributed between the two interaction clusters.

### UV cross-linking in the frozen state indicates improved reaction efficiency

Importantly, the first detected interaction cluster within FOX1-RRM reveals Pro125 as the dominant residue involved in the FBE interaction, whereas previous studies have reported the adjacent Phe126 as the main contact site.^10,22^ We therefore sought to pinpoint the interacting protein residue using the targeted approach and similar data analysis logic to that applied to TDP-43 complexes. As a result, we confirmed the Phe126 and Phe160 residues as the main interaction sites within FOX1-RRM, while Pro125 was a result of the initial misassignment of the DDA workflow. Interestingly, our targeted approach reveals a secondary interaction site in the Phe126 cluster, His120. Yet, His120 is confidently observed only when cross-linked to U nucleobase in the AU RNA composition, the same as reported by NMR as a His120–U5 interaction.^22^ Our data suggests that the FOX1-RRM–FBE interaction is predominantly driven by Phe residues, while His120-based contacts are only revealed in the context of interactions with U5 nucleotide in FBE.

To directly compare the efficiency of UV cross-linking across temperatures during sample irradiation, we applied our PRM-based approach to perform label-free quantification at MS2 level of selected cross-links, mapping to two individual interaction clusters (Fig. 4E). We observed that both cross-links, regardless of the interaction site they map to, were more abundant in samples irradiated in the frozen state. This was particularly pronounced in the Phe126 cluster, where the cross-link to G nucleobase in the CGU RNA composition showed up to a 5-fold increase in abundance relative to samples held at non-negative temperatures during irradiation. However, it is unclear whether the beneficial effect of the frozen state on the UV cross-linking efficiency can be generalized. Therefore, it remains to be investigated how conformational and physicochemical properties of peptide–RNA cross-links correlate with sample temperature during UV irradiation.

## Discussion

Protein–RNA complexes play a crucial role in the regulation of many fundamental biological processes. For a comprehensive understanding of these processes, it is necessary to understand how proteins and RNA interact. UV XL-MS is a widely used approach for obtaining structural insights into protein–RNA interactions. CLIR-MS, as one of the protein–RNA XL-MS workflows, allows the characterization of these interactions at site-specific resolution. It has been successfully used to generate distance constraints for integrative structure modeling of protein–RNA complexes, or to uncover novel RNA interaction sites in RBPs. However, CLIR-MS, like many other DDA-based methods in the field, is not designed for comparative quantification of protein–RNA interactions.

The introduction of our PRM-based workflow enables more reproducible and sensitive detection of peptide–RNA cross-links present in low abundance. It also improves the structural characterization of protein–RNA interactions by increasing the confidence of site localization on both protein and RNA components. Importantly, our novel approach enables the comparative label-free quantification of changes in abundance of cross-linked RNA, resulting from complex conformational changes, PTM-driven modulation of interaction or protein mutations and truncations. To our knowledge, PRM-based acquisition has not been applied to peptide–RNA cross-links, although selected/multiple reaction monitoring has been used in the past.^47^ Combined with DDA-based CLIR-MS, this workflow enables both qualitative and quantitative characterization of protein–RNA interactions.

Our presented approach was applied to an *in vitro* model system, the NTD of TDP-43 protein, complexed with specifically recognized (GU)_6_ RNA. In the qualitative analysis, we reveal two RNA interaction sites, Cys39 and Cys50 located in the oligomerization interface of the protein. While our previous MS experiments already revealed Cys50 as a novel RNA interaction site of TDP-43, Cys39 has not been reported before in this context.^8^ Interestingly, Cys39 is located outside of the head and tail regions, in the α1-β3 loop of TDP-43 NTD, recognized as the M1 mitochondrial targeting sequence (MTS, 35-FPGACGL-41). Deletion of the M1 sequence has been shown to greatly reduce TDP-43 mitochondrial accumulation, linked to proteinopathy in ALS and FTD.^48^ Remarkably, high-affinity RNA interaction sites (Phe147, Phe149) and other MTS, M3, are shared in the RRM1 domain of TDP-43 (146-GFGFV-150), making the recognition of the mitochondrial target sequence possibly highly dependent on RNA binding.^49^ This poses an intriguing question of probable functional interplay between RNA interaction, oligomerization and mitochondrial localization of TDP-43, all coordinated by the Cys39 residue.

By uncovering Cys39 and Cys50 as novel RNA cross-linking sites, we demonstrate that NTD can interact with RNA independent of high-affinity RRM domains and that oligomerization and RNA interactions are shared by the same protein region, strengthening our previous finding.^8^ By including the cross-link-specific and RNA-specific fragment ions in the spectra annotation procedure, we determine U base as the covalently bound remnant on the peptide chain, and 5′-GU-3′ as the dominant RNA sequence bound. These observations are shared between Cys39 and Cys50.

Cysteine–RNA UV cross-links are well documented in the literature, with the first structurally characterized cysteine-uracil photoproduct dating back to the 1960s.^50^ Cysteine is one of the most photoreactive amino acids, owing to the high reactivity of the thiyl radical.^51,52^ Unlike aromatic residues, cysteine does not require π-stacking interactions with nucleobases to produce a stable, highly reactive radical.^10^ Consequently, cysteine may get cross-linked to a nucleobase even when the protein–RNA interaction is transient – providing evidence of spatial proximity, but not necessarily of a stable, functionally essential molecular contact.^53^ In our case, however, UV cross-linking was performed on pre-incubated TDP-43 NTD–(GU)_6_, allowing formation of stable protein–RNA complexes before irradiation. The detected cross-links therefore reflect specific contact sites rather than incidental ones.

Our previous MS and molecular dynamics data indicated Cys-driven RNA contacts, whereas NMR instead identified arginine as the key interaction contributor.^8^ The majority of residues detected by NMR lie within the oligomerization interface, and a contact mediated by any of these could be then structurally plausible. Collectively, our findings indicate that (GU)_6_ binding is governed cooperatively by the oligomerization interface as a whole, whereas each method preferentially reports the subset of contacts to which it is chemically most sensitive — in the case of UV cross-linking, the photochemically most reactive residues. The protein–RNA interaction sites identified by the different approaches are therefore complementary rather than mutually exclusive.

As a part of the quantitative analysis, we observe enhanced RNA interactions in the oligomerization-deficient NTD Tm mutant, compared to the oligomeric NTD WT. Both interaction sites show a comparable degree of response to RNA interactions upon protein monomerization, despite varying surface exposure. Moreover, we show that titration of (GU)_6_ induces the dissociation of NTD WT oligomers, suggesting that the multivalent interactions driven by 12-mer RNA sequence are sufficient to outcompete weaker, predominantly electrostatic interactions facilitating protein oligomerization. We also demonstrate that surface-exposed Cys39 engages RNA more effectively than the Cys50 residue that is buried in the oligomerization interface.

We detect consistent differences in chromatographic and titration behavior of two RNA forms of the same peptide that differ by a single phosphate group. The origin of these observations remains to be uncovered. Nevertheless, we hypothesize that one possible explanation of the presence of an additional phosphate group at the 3′ end of GU dinucleotide) lies in the arginine-phosphate group interaction (arginine fork),^54,55^ where an intramolecular guanidinium–phosphate salt bridge could shield the 3′-end phosphate from the dephosphorylation step, thus yielding the phosphate-retaining form of GU dinucleotide. Supporting this hypothesis, the phosphate-retaining form is observed only when the cross-linked peptide contains at least one Arg residue. These Arg residues are located near either the peptide N terminus of Cys50-containing peptides or the C terminus of Cys39-containing peptides. Their terminal positions may provide sufficient conformational flexibility for formation of the proposed arginine forks.

Yet, it is difficult to extend this hypothesis to the initial protein–RNA complex formation step. A potential arginine fork may be formed between one of the Arg residues and the phosphate group of a phosphodiester bond, simultaneously with the cross-link formation between Cys and U base. This dual interaction within a single peptide–RNA cross-linking interface could cooperatively stabilize the interaction^10,56^ but is less favorable to obtain because it imposes more geometric constraints. This could explain the higher inflection points, consistently observed for the phosphate-retaining form of GU dinucleotide.

We additionally hypothesize that the stronger retention of the phosphate-retaining form results from a conformational effect, in which the additional phosphate group increases the exposure of the hydrophobic regions of peptide–RNA cross-links. Assuming this effect, separation and retention would be governed by this proposed conformational change rather than by the polarity of the added single phosphate group alone. This behavior could suggest the involvement of arginine-driven interactions: the same arginine fork could act as an intramolecular ion-pairing agent that counteracts the expected polarity-driven shift toward earlier elution. Hypothesized interaction appears geometrically and electrostatically plausible during the experimental steps considered (UV cross-linking, 3′ dephosphorylation, RP-HPLC separation), although its presence at each step remains to be demonstrated.

Collectively, we provide evidence that TDP-43 NTD oligomerization and RNA interaction are mutually exclusive processes, with RNA possessing abilities to trigger protein monomerization. Given the importance of NTD-driven protein oligomerization in TDP-43 phase separation,^31,33^ we speculate that the monomerization process induced by the presence of RNA may serve as a regulatory mechanism for the formation of biomolecular condensates by the full-length TDP-43. As shown in the example of the phosphomimetic S48E variant of the TDP-43 NTD, introduction of a single negative charge at the oligomerization interface reduces both protein oligomerization and LLPS, which demonstrates that the oligomeric state of protein is a key determinant for phase separation.^33^ It is therefore plausible that the presence of RNA would similarly reduce LLPS by shifting the oligomerization equilibrium towards monomeric form. However, the observed effect of RNA on the NTD monomerization remains to be tested in the full-length protein, particularly given the distinct contributions of the RRMs to RNA binding and the C-terminal domain (CTD) to LLPS.

Moreover, specific RNA binding to GU-rich sequences has been shown before to prevent full-length TDP-43 oligomerization and subsequential aggregation, emphasizing the contribution of RRM1 and LCD domains, but not the NTD.^57,58^ Therefore, NTD-driven RNA binding might serve as a regulatory mechanism of maintaining TDP-43 oligomerization.

We also hypothesize that RNA interactions driven by the NTD might contribute to specific recognition of RNA sequences and their distinction from linear single-stranded DNA (ssDNA), rich in TG-repeats. TDP-43 can interact with both types of nucleic acids. Interestingly, it uses the same residues within RRM1 (Trp113, Phe194) and RRM2 (Lys263, Arg227) domains to bind both linear single-stranded RNA (ssRNA) and linear ssDNA, which are rich in UG-and TG-repeats, respectively. Within RRM1, the same stacking interactions drive the coordination of U and T. The differential methyl group on T is not recognized by any TDP-43 residue. Within RRM2, the RNA-specific contacts incorporate the 2′-OH group; however, when bound to ssDNA, the same amino acid side chains interact with the ssDNA phosphodiester backbone.^59^ As the functional outcome of interaction with each type of nucleic acid is highly diverse, it might be necessary to employ different protein domains or regions to ensure specificity of recognition. The cooperativity between canonical RNA-binding domains and the NTD may therefore be vital for specific RNA recognition and its distinction from ssDNA, especially when both nucleic acids lack defined higher-order structure.

We also successfully applied our combined workflow to investigate temperature-dependent differences in the efficiency of cross-link formation, using the FOX1-RRM–FBE complex as a model system. UV cross-linking at temperatures higher than the conventionally used ∼0 °C has been reported to significantly increase the photoreactivity of several proteins, whereas lower temperatures proved beneficial for reaction selectivity.^60^ Our qualitative analysis revealed minor differences in the total number of cross-links detected across conditions; however, we observed differences while comparing the number of cross-links detected between the main interaction clusters. Selected cross-links mapping to both clusters showed more efficient cross-link formation in samples present in the frozen state during UV irradiation. Sample freezing could lock the relative orientation of interacting amino acid side chains and nucleobases and thereby increase the yield of the cross-linking reaction while also slowing competing photodamage. Yet the reaction efficiency in the frozen state could be impaired if the minor conformational adjustments needed to reach the photoreactive geometry are restricted by the loss of atomic fluctuations, especially in the context of transient interactions.^61^ Further, more systematic investigation and optimization of UV cross-linking in the frozen state could reveal its potential, despite several limitations related to sample handling in this condition, such as maintaining low temperature throughout the process or potential UV light scattering.

Collectively, we demonstrate that the specific RNA recognition by FOX1-RRM is not affected by varying temperature during the UV cross-linking procedure, as shown by the consistent identification of the two dominant cross-linking sites, clustering around Phe126 and Phe160. The temperature-independent FBE recognition is likely determined by the remarkably high affinity of complex formation.^22^ Our procedure offers a valuable approach for exploring the mechanisms of cross-link formation in the context of temperature-controlled conformational stability of a protein–RNA complex. Moreover, it can be used for exhaustive optimization of UV cross-linking protocols, allowing to compare the effect of other factors, such as buffers, photosensitizers, or protein–RNA complex incubation time, on the efficiency of cross-link formation.

While our workflow supports comparative analysis of peptide–RNA cross-link abundance across selected conditions, it is not designed to deliver absolute affinities or kinetics of the targeted interactions. Cross-link formation irreversibly traps interaction-engaged residues through covalent bonds, thereby perturbing the reversible association-dissociation equilibrium from which affinity and reaction rate constants could be otherwise derived. Moreover, “zero-length” UV cross-linking yield often reflects photochemical reactivity rather than binding strength, since it is dictated by the local geometry of the interacting and adjacent residues, as well as the amino acid and nucleotide composition in the interaction hotspot.^7,10,62,63^ Finally, because potentially any amino acid may form a cross-link, the combinatorial search space considerably expands and complicates the accurate identification of a cross-link by increasing the number of false-positive identifications so that the score threshold needs to be raised to achieve the same statistical significance.^64–66^ However, a better understanding of concurrent photoreactions, resulting in variety of photochemical products, could improve both the yield and the detection confidence of peptide–RNA cross-links.

Collectively, our framework provides a comprehensive comparison of protein–RNA interactions across structural and biophysical conditions. In combination with complementary techniques, our improved structural readout could provide spatial constraints for integrative structural modeling of protein–RNA complexes, as well as elucidate the molecular recognition of target RNA by proteins. Quantitative analysis of peptide–RNA abundance, particularly in the context of the titration experiment, could further reveal how RNA binding reshapes protein conformation, offering insight into the interplay between complex structure and RNA interaction.

## Materials and methods

### Protein expression and purification

Recombinant 6xHis-tagged TDP-43 NTD WT and TDP-43 NTD Tm were prepared as described previously.^8^ Both protein constructs were overexpressed in One Shot BL21 Star (DE3) Escherichia coli cells (Thermo Fisher Scientific). Cells were grown in Luria-Bertani (LB) broth base (Invitrogen) for 30 h at 25 °C after induction at an optical density of 1.0 with 1 mM isopropyl-β-D-thiogalactopyranoside (IPTG, Biosolve). Cells were harvested by centrifugation and resuspended in 20 mL lysis buffer (50 mM sodium phosphate buffer pH 8.0, 300 mM sodium chloride (NaCl), 10 mM imidazole (all Sigma-Aldrich), 1 mM 1,4-dithiothreitol (DTT, Thermo Fisher Scientific), supplemented with 2 protease inhibitor tablets (Roche) per 1 L culture. The cell suspension was homogenized using a W-385 LSL Secfroid rod sonicator (Heat System Ultrasonic) with 11 cycles (10 s on, 20 s off) and subsequently centrifuged (Sorvall SS-34 rotor, 15,000 × g, 4 °C, 20 min). The supernatant was filtered through 0.22 μm syringe filters and applied to pre-equilibrated Ni Sepharose Excel material (Sigma-Aldrich) packed in a gravity Econo-Column (Biorad).

The column was washed once with wash buffer 1 (50 mM sodium phosphate buffer pH 8.0, 1 M NaCl, 10 mM imidazole, 1 mM DTT) and a second time with wash buffer 2 (50 mM sodium phosphate buffer pH 8.0, 1 M NaCl, 30 mM imidazole, 1 mM DTT). Protein was eluted in 30 mL elution buffer (50 mM sodium phosphate buffer pH 8.0, 1 M NaCl, 500 mM imidazole, 1 mM DTT) and dialyzed overnight in dialysis buffer (5 mM sodium phosphate buffer pH 7.4, 1 mM NaCl, 1 mM DTT). Protein concentration was determined by the extinction coefficients and molecular weights of the constructs (ExPASy ProtParam software)^67^ and absorbance at 280 nm (NanoDrop, Thermo Scientific). Protein was concentrated using a Vivaspin 5,000 MWCO concentrator (Sartorius Stedim Biotech), aliquoted, flash frozen, and stored at-80 °C.

Recombinant 6xHis-tagged FOX1-RRM construct was overexpressed in BL21 (DE3) Escherichia coli cells (Thermo Fisher Scientific). Cells were grown in LB broth base (Invitrogen) for 4 h at 37 °C after induction at an optical density of 0.6 with 1 mM IPTG. Cells were harvested by centrifugation and resuspended in 20 mL lysis buffer (50 mM sodium phosphate buffer pH 8.0, 300 mM NaCl, 10 mM imidazole), supplemented with 0.002% (v/v) SUPERase RNase inhibitor (Ambion) per 1 L culture. The cell suspension was lysed by two passes through a cell cracker (Avestin) and subsequently centrifuged at 20,000 × g, 4 °C, 20 min. The supernatant was applied to pre-equilibrated Ni-NTA (nickel-nitrilotriacetic acid, Qiagen) resin packed in a gravity Econo-Column (Biorad). After washing with lysis buffer, the protein was eluted with a step gradient of imidazole (20–500 mM). Eluted fractions of highest purity, as judged by SDS-PAGE assay, were subjected to a second Ni-NTA affinity purification procedure. Eluted protein was dialyzed overnight in dialysis buffer (10 mM sodium phosphate buffer pH 6.5, 20 mM NaCl). Protein concentration was determined by the extinction coefficients and molecular weights of the constructs (ExPASy ProtParam software)^67^ and absorbance at 280 nm (NanoDrop, Thermo Scientific). Protein was concentrated using a Vivaspin 5,000 MWCO concentrator (Sartorius Stedim Biotech), aliquoted, flash frozen, and stored at-80 °C.

### RNA oligonucleotides

(GU)_6_ RNA (5′-GUGUGUGUGUGU-3′) and FBE (5′-UGCAUGU-3′) were purchased from IDT. Lyophilized RNA samples were resuspended in RNase-free water to a final concentration of 1.4 mM, aliquoted, and stored at-80 °C.

### Protein–RNA UV cross-linking

For the initial interaction site discovery experiment, a reaction volume of 200 μL was prepared in biological quadruplicates for both protein variants, containing a final concentration of 50 μM (total amount of 100 μg) TDP-43 NTD WT (or TDP-43 NTD Tm) and a final concentration of 50 μM isotopically unlabeled (GU)_6_ RNA, in 5 mM sodium phosphate buffer pH 7.4, 1 mM NaCl, 1 mM DTT. After 30 min of incubation on ice, samples were irradiated on ice at 254 nm and at total energy of 3.2 J/cm^2^ (in 0.8 J/cm^2^ increments) on a 48-well plate (Spectrolinker XL-1500 UV Crosslinker, Spectronics Corporation). Each irradiation step was followed by a pause of 1 min to allow the sample to cool. The titration experiment was performed as described above, except for varied (GU)_6_–TDP-43 NTD WT molar ratios, ranging from 0 to 2.5 (at fixed final protein concentration of 50 μM and using a total amount of 100 μg of protein).

The FOX1 experiment was performed in a reaction volume of 500 μL in biological quadruplicates. Samples contained a final concentration of 15 μM (total amount of 100 μg) FOX1-RRM and a final concentration of 15 μM isotopically unlabeled FBE RNA (total amount of 16 μg), in 10 mM sodium phosphate buffer pH 6.5, 50 mM NaCl. The FOX1-RRM–FBE complex was *in vitro* reconstituted on ice and subsequently incubated for 20 min at either-80 °C,-20 °C, 0 °C, RT or 37 °C. Samples were irradiated at 254 nm and at total energy of 3.2 J/cm^2^ (in 0.8 J/cm^2^ increments) on a 12-well plate (Spectrolinker XL-1500 UV Crosslinker, Spectronics Corporation). Each irradiation step was followed by a pause of 1 min to allow the sample to cool. Reaction temperature was maintained either by filling up the plate holder with dry or wet ice, or with heated water.

Samples from both experiments were subsequently processed based on the same protocol, described previously.^11,14^ Samples were precipitated overnight using ethanol and sodium acetate. Dried pellets were resuspended in 50 mM Tris-HCl pH 7.9, 4 M urea) and further diluted with 50 mM Tris-HCl pH 7.9, to a final concentration of 1 M urea. Samples were digested with a mixture of RNase A (Roche) and RNase T1 (Thermo Scientific) for 2 h at 52 °C (5 ug RNase A and 5 U RNase T1 per mg of protein–RNA complex), followed by digestion with chymotrypsin (Roche) for 2 h at 25 °C, at a 1:50 enzyme-protein ratio (w/w). The reaction was inactivated with formic acid (2% v/v). Samples were cleaned up by solid-phase extraction (SPE, SepPak 50 mg tC18 cartridges, Waters), and dried by vacuum centrifugation. Resuspended pellets were subject to ^18^O labeling,^6^ using a 1:1 mixture of light ATP (Sigma-Aldrich) and heavy ^18^O_4_-γ-ATP (synthesized as described below). The labeling reaction was carried out for 1 h at 37 °C in LoBind tubes (Eppendorf), followed by a second SPE clean-up. Enrichment of protein– RNA cross-links was performed using metal oxide affinity chromatography (MOAC), followed by STAGE tip C18 SPE using self-packed tips.^6,12^ Samples were dried by vacuum centrifugation.

### Liquid chromatography–tandem mass spectrometry (LC-MS/MS)

Samples from both experiments were resuspended in 15 μL of water/acetonitrile/formic acid (95:5:0.1, v/v/v).

For the data-dependent acquisition-based (DDA) CLIR-MS workflow, 10% volume of each replicate per protein–RNA complex was pooled and 1 μL was used for analysis in technical duplicates. LC-MS/MS analysis was performed with an Easy nLC-1200 HPLC system coupled to an Orbitrap Fusion Lumos mass spectrometer equipped with a Nanoflex nanoflow electrospray ionization source (all Thermo Fisher Scientific). Samples were separated on an Acclaim PepMap RSLC C18 column (250 mm × 75 µm i.d., 2 μm particle size, Thermo Fisher Scientific) at a constant flow rate of 300 nL/min, with mobile phases A = water/acetonitrile/formic acid (98:2:0.15, v/v/v) and B = acetonitrile/water/formic acid (80:20:0.15, v/v/v). A linear 6–40% B gradient within 60 min was used for the separation. The Orbitrap Fusion Lumos was used in the positive mode. Precursor ion spectra were acquired in the Orbitrap analyzer at a resolution of 120,000 in 3 s cycles, with a scan range of 350–1500 m/z. Fragmentation of selected precursors (2–7^+^ charge state, 1.2 m/z isolation window) was achieved with higher-energy collisional dissociation (HCD), using a normalized stepped collision energy of 23 ± 5%. Dynamic exclusion was activated for 30 s after one sequencing event.

For quantitative parallel reaction monitoring (PRM), single injections of 1 μL (TDP-43 NTD variants comparison experiment, FOX1-RRM experiment) and 4 μL (TDP-43 NTD WT–(GU)_6_ titration experiment) of each replicate were performed. Chromatographic separation was performed as described above.

Target precursor ions of peptide–RNA cross-links were selected upon analysis of the CLIR-MS data, using the following criteria: presence of the precursor ion in all experimental conditions, precursor ion tolerance = 10 ppm, apex retention time more than 10 min into the gradient. Excluded were precursor ions characterized by broad chromatographic elution peaks of irregular profiles and without clearly defined peak apex. Target precursor ions were specified as a list of peptide–RNA cross-links with different neutral losses, defined by their m/z ratios and present in two charge states (2^+^, 3^+^), resulting from both ^16^O labeling (light state) and ^18^O labeling (heavy state). Precursor ion spectra of desired targets (Suppl. Tab. 1) were acquired in the Orbitrap analyzer at a resolution of 60,000 in a retention time-scheduled manner. Wide retention time windows were specified owing to the broad elution profiles of the target cross-links and their appearance as several chromatographic peaks. Targeted fragmentation was achieved with HCD, using normalized stepped collision energy of 23 ± 5%. Fragment ions were detected in the Orbitrap at a resolution of 60,000, with a scan range of 100–2000 m/z, an AGC target of 4.0e5, a maximum injection time of 200 ms and maximum cycle times of 2.9 s (TDP-43 NTD variants comparison experiment), 2.5 s (TDP-43 NTD WT– (GU)_6_ titration experiment) and 5.0 s (FOX1-RRM experiment).

## Data analysis

For DDA-based CLIR-MS data analysis, MS files were converted from the.raw to centroided.mzXML file fomat using msconvert.exe (ProteoWizard msConvert v.3.0.9393c).^68^ Converted files were searched using xQuest (version 2.1.53, available from https://gitlab.ethz.ch/leitner_lab/xquest_xprophet)^69,70^ against a database containing FASTA sequences of TDP-43 NTD protein variants or FOX1-RRM protein, their decoy sequences, and RNA sequences (either (GU)_6_ in both TDP-43 NTD experiments or FBE in the FOX1-RRM experiment). The reverse decoy protein database was generated using the *xdecoy.pl* script included with xQuest. No contaminant protein sequences have been included in the search due to their negligible abundances. The maximum length of an RNA adduct was set to four nucleotides for all experiments. Oxidation of methionine was set as a variable protein modification. MS spectra were paired according to the mass shift of 6.012735 Da resulting from ^18^O labeling. Additional search settings were: enzyme = chymotrypsin (FWYML/P) in the TDP-43 NTD experiments or trypsin (KR/P) in the FOX1-RRM experiment, the maximum number of missed cleavages = 4 in the TDP-43 NTD experiment or 2 in the FOX1-RRM experiment, precursor ion tolerance = 10 ppm, fragment ion tolerance = 15 ppm. Identifications from xQuest were post-processed using the RNxQuest 1.0.0 Python package.^64^ The false discovery rate was set to 1% at the peptide–RNA conjugate level.

PRM-based quantification of target cross-links was performed on.raw MS measurement files using Skyline software (64-bit, 22.2.0.351).^71^ Transition lists containing all possible unmodified (non-cross-linked, “common”) and modified (cross-linked, “XL”) peptide backbone and RNA backbone fragments were generated using custom Python scripts. Precursor ions were manually identified based on the co-elution of light and heavy fragment ions, matching the intact peak shape. Peak boundaries were adjusted to exclude interfering and/or closely eluting peaks. The abundance of each cross-link was calculated by summing up the integrated peak area of the five most intense fragment ions per precursor in the light channel, prioritizing cross-linked fragments. Obtained values were normalized to the peak area of a non-cross-linked TDP-43 NTD peptide (69-GNLVY-73), located outside of the oligomerization interface and chromatographically resolved from all analyzed cross-links. In the TDP-43 NTD variants comparison experiment, the significance of abundance changes was assessed using a paired *t*-test, with p-values adjusted for multiple comparisons by the Benjamini–Hochberg procedure. Statistical significance was denoted as follows: * for q < 0.05, ** for q < 0.01, *** for q < 0.001. Abundance change was represented as mean ± standard error mean (SEM) of n = 4, per cross-link. In the titration experiment, datapoints representing normalized peak areas measured across RNA-protein molar ratios were fitted to a three-parameter equation. Replicate 2 at 2.5 RNA-protein ratio was excluded due to its relative normalization peak area falling outside the 95% confidence interval of the distribution across all measurements. The corresponding cross-link abundance values were affected by this normalization error and were not analyzed. This yielded n = 4 replicates per measured cross-link abundance for all RNA-protein ratios except for RNA-protein ratio of 2.5 (n = 3). The 95% confidence band around each fit represents mean predicted response. In the FOX1-RRM experiment, obtained peak integration values of the selected five fragment ions were normalized to the peak area of a non-cross-linked FOX1-RRM peptide (135-QMFGQFGK-142), located outside of the RNA interaction interface and chromatographically resolved from all analyzed cross-links. Abundance change was represented as mean ± standard error mean (SEM) of n = 4, per cross-link. Statistical tests and curve fitting were performed with JMP Student software (19.0.4; JMP Statistical Discovery LLC, Cary, NC).

### Heavy ^18^O_4_-γ-ATP synthesis

ATP-(^18^O)_4_ trisodium salt was synthesized following the previously published methods^72,73^ with the following major modifications. Both the unlabeled potassium dihydrogen phosphate (99.5% pure, Fluka #60220) and the labeled potassium dihydrogen phosphate (¹8O₄, 95% isotopic purity, 97% chemical purity, Cambridge Isotope Laboratories #OLM-7493-1) were purchased commercially. The alkali metal salts of ADP and dihydrogen phosphate were converted into their respective ammonium salts using Dowex 50W X8 (200–400 mesh, Supelco #44519) instead of Dowex 50W X8 (100–200 mesh). To form the ammonium counterions of ADP and dihydrogen phosphate, tri-n-butylamine (99% pure, Sigma Aldrich #90780) was used in addition to tri-n-ethylamine (99% pure, Sigma Aldrich #8.08352) and tri-n-propylamine (98% pure, AlfaAesar #A10285.AK). The ADP imidazolate was precipitated as a sodium salt prior to the subsequent reaction to form ATP, and a reverse-phase chromatographic system (Biotage Isolera One equipped with a FlashPure IC C18 column (Büchi #145152991)) was used for purification instead of size exclusion chromatography. The synthesis scheme is provided in the *Supplementary Information* section (Suppl. Fig. 3).

### Native polyacrylamide gel electrophoresis (PAGE)

10 μg of each UV cross-linked TDP-43 NTD WT–(GU)_6_ complex, reconstituted at varying RNA-to-protein ratios, were mixed with Novex Tris-Glycine Native Sample Buffer (2×, Invitrogen). Samples were loaded onto 16% Novex Tris-Glycine Mini Protein Gels (Invitrogen). Electrophoresis was carried out in ice-cold Novex Tris-Glycine Native Running Buffer (1×) at 200 V for 40 min. Gels were stained with SimplyBlue SafeStain (all Invitrogen) according to the manufacturer’s protocol. NativeMark Unstained Protein Standard (Invitrogen) was used for estimating the molecular weight of separated molecules.

### Structural image representation

Figures displaying protein structures were generated using PyMOL graphic software (version 3.1.6.1, Schrodinger, LLC).

## Acknowledgements

A.L. would like to thank Ruedi Aebersold and Paola Picotti for access to instrumentation and laboratory infrastructure.

## Declaration of AI-assisted technologies in the writing process

During the preparation of this work, the authors used the OpenAI GPT-6 model to improve the grammar and clarity of the *Introduction* section and to assist in refining custom Python scripts for generating transition lists imported into Skyline software. The authors reviewed and verified the suggestions and take full responsibility for the published content.

## Author contributions

I.S. and A.L. developed the main research idea, overall study design, and designed the methodology used in this study. A.L., M.P., J.H., F.H-T.A. and A.Z. provided scientific advice and guidance for the study. A.Z. and J.J. performed sample preparation by overexpressing and purifying recombinant proteins. J.O. carried out the synthesis of the heavy ATP isotopolog. I.S. performed sample preparation for mass spectrometry (MS), analyzed the data and visualized the results. I.S. prepared the manuscript figures and schematics and wrote the original draft of the manuscript. I.S. and A.L. reviewed and revised the manuscript. A.L., and F.H-T.A. acquired funding and supervised the project.

## References

1. M. W. Hentze, A. Castello, T. Schwarzl, T. Preiss, A brave new world of RNA-binding proteins. Nat. Rev. Mol. Cell Biol. 19, 327–341 (2018). doi:10.1038/nrm.2017.130.

2. P. Wu, Inhibition of RNA-binding proteins with small molecules. Nat. Rev. Chem. 4, 441–458 (2020). doi:10.1038/s41570-020-0201-4.

3. A. R. Julio, K. M. Backus, New approaches to target RNA binding proteins. Curr. Opin. Chem. Biol. 62, 13–23 (2021). doi:10.1016/j.cbpa.2020.12.006.

4. R. M. L. Queiroz, T. Smith, E. Villanueva, M. Marti-Solano, M. Monti, M. Pizzinga, D.-M. Mirea, M. Ramakrishna, R. F. Harvey, V. Dezi, G. H. Thomas, A. E. Willis, K. S. Lilley, Comprehensive identification of RNA–protein interactions in any organism using orthogonal organic phase separation (OOPS). Nat. Biotechnol. 37, 169–178 (2019). doi:10.1038/s41587-018-0001-2.

5. J. Trendel, T. Schwarzl, R. Horos, A. Prakash, A. Bateman, M. W. Hentze, J. Krijgsveld, The human RNA-binding proteome and its dynamics during translational arrest. Cell 176, 391–403 (2019). doi:10.1016/j.cell.2018.11.004.

6. M. Götze, C. P. Sarnowski, T. de Vries, A. Knörlein, F. H.-T. Allain, J. Hall, R. Aebersold, A. Leitner, Single nucleotide resolution RNA–protein cross-linking mass spectrometry: A simple extension of the CLIR-MS workflow. Anal. Chem. 93, 14626–14634 (2021). doi:10.1021/acs.analchem.1c02384.

7. K. Kramer, T. Sachsenberg, B. M. Beckmann, S. Qamar, K.-L. Boon, M. W. Hentze, O. Kohlbacher, H. Urlaub, Photo-cross-linking and high-resolution mass spectrometry for assignment of RNA-binding sites in RNA-binding proteins. Nat. Methods 11, 1064–1070 (2014). doi:10.1038/nmeth.3092.

8. A. Zbinden, I. Smok, F. Pellegrini, A. Oertel, A. Cléry, V. I. Wiersma, W.-P. Huang, B. Gatta, W. Zhong, Y. Li, M. Götze, M. Peter, N. Bargenda, M. Hruska-Plochan, A. Caflisch, P. Fratta, T. A. Shelkovnikova, F. H.-T. Allain, A. Leitner, M. Polymenidou, Monomeric TDP-43 binds GU-rich RNA via its NTD but only its oligomers block cryptic exon inclusion. bioRxiv (2026). doi:10.64898/2026.10.01.755875.

9. E. Kühn-Hölsken, C. Lenz, A. Dickmanns, H.-H. Hsiao, F. M. Richter, B. Kastner, R. Ficner, H. Urlaub, Mapping the binding site of snurportin 1 on native U1 snRNP by cross-linking and mass spectrometry. Nucleic Acids Res. 38, 5581–5593 (2010). doi:10.1093/nar/gkq272.

10. A. Knörlein, C. P. Sarnowski, T. de Vries, M. Stoltz, M. Götze, R. Aebersold, F. H.-T. Allain, A. Leitner, J. Hall, Nucleotide-amino acid π-stacking interactions initiate photo cross-linking in RNA-protein complexes. Nat. Commun. 13, 2719 (2022). doi:10.1038/s41467-022-30284-w.

11. G. Dorn, A. Leitner, J. Boudet, S. Campagne, C. von Schroetter, A. Moursy, R. Aebersold, F. H.-T. Allain, Structural modeling of protein–RNA complexes using crosslinking of segmentally isotope-labeled RNA and MS/MS. Nat. Methods 14, 487–490 (2017). doi:10.1038/nmeth.4235.

12. C. P. Sarnowski, A. Knörlein, T. de Vries, M. Götze, I. Beusch, R. Aebersold, F. H.-T. Allain, J. Hall, A. Leitner, A highly sensitive protein-RNA cross-linking mass spectrometry workflow with enhanced structural modeling potential. Nucleic Acids Res. 53, gkaf523 (2025). doi:10.1093/nar/gkaf523.

13. G. Dorn, C. Gmeiner, T. de Vries, E. Dedic, M. Novakovic, F. F. Damberger, C. Maris, E. Finol, C. P. Sarnowski, J. Kohlbrecher, T. J. Welsh, S. Bolisetty, R. Mezzenga, R. Aebersold, A. Leitner, M. Yulikov, G. Jeschke, F. H.-T. Allain, Integrative solution structure of PTBP1-IRES complex reveals strong compaction and ordering with residual conformational flexibility. Nat. Commun. 14, 6429 (2023). doi:10.1038/s41467-023-42012-z.

14. T. de Vries, M. Novakovic, Y. Ni, I. Smok, C. Inghelram, M. Bikaki, C. P. Sarnowski, Y. Han, L. Emmanouilidis, G. Padroni, A. Leitner, F. H.-T. Allain, Specific protein-RNA interactions are mostly preserved in biomolecular condensates. Sci. Adv. 10, eadm7435 (2024). doi:10.1126/sciadv.adm7435.

15. P. I. Koukos, A. M. J. J. Bonvin, Integrative modelling of biomolecular complexes. J. Mol. Biol. 432, 2861–2881 (2020). doi:10.1016/j.jmb.2019.11.009.

16. K. Barkovits, S. Pacharra, K. Pfeiffer, S. Steinbach, M. Eisenacher, K. Marcus, J. Uszkoreit, Reproducibility, specificity and accuracy of relative quantification using spectral library-based data-independent acquisition. Mol. Cell. Proteomics 19, 181–197 (2020). doi:10.1074/mcp.RA119.001714.

17. A. C. Peterson, J. D. Russell, D. J. Bailey, M. S. Westphall, J. J. Coon, Parallel reaction monitoring for high resolution and high mass accuracy quantitative, targeted proteomics. Mol. Cell. Proteomics 11, 1475–1488 (2012). doi:10.1074/mcp.O112.020131.

18. N. Rauniyar, Parallel reaction monitoring: A targeted experiment performed using high resolution and high mass accuracy mass spectrometry. Int. J. Mol. Sci. 16, 28566–28581 (2015). doi:10.3390/ijms161226120.

19. A. Bourmaud, S. Gallien, B. Domon, Parallel reaction monitoring using quadrupole-Orbitrap mass spectrometer: Principle and applications. Proteomics 16, 2146–2159 (2016). doi:10.1002/pmic.201500543.

20. C. Iacobucci, C. Piotrowski, R. Aebersold, B. C. Amaral, P. Andrews, C. Haupt, C. Sailer, F. Stengel, A. Leitner, A. Sinz, First community-wide, comparative cross-linking mass spectrometry study. Anal. Chem. 91, 6953–6961 (2019). doi:10.1021/acs.analchem.9b00658.

21. P. J. Lukavsky, D. Daujotyte, J. R. Tollervey, J. Ule, C. Stuani, E. Buratti, F. E. Baralle, F. F. Damberger, F. H.-T. Allain, Molecular basis of UG-rich RNA recognition by the human splicing factor TDP-43. Nat. Struct. Mol. Biol. 20, 1443–1449 (2013). doi:10.1038/nsmb.2698.

22. S. D. Auweter, R. Fasan, L. Reymond, J. G. Underwood, D. L. Black, S. Pitsch, F. H.-T. Allain, Molecular basis of RNA recognition by the human alternative splicing factor Fox-1. EMBO J. 25, 163–173 (2006). doi:10.1038/sj.emboj.7600918.

23. E. Buratti, F. E. Baralle, Characterization and functional implications of the RNA binding properties of nuclear factor TDP-43, a novel splicing regulator of CFTR exon 9. J. Biol. Chem. 276, 36337– 36343 (2001). doi:10.1074/jbc.M104236200.

24. S. H. Ou, F. Wu, D. Harrich, L. F. García-Martínez, R. B. Gaynor, Cloning and characterization of a novel cellular protein, TDP-43, that binds to human immunodeficiency virus type 1 TAR DNA sequence motifs. J. Virol. 69, 3584–3596 (1995). doi:10.1128/JVI.69.6.3584-3596.1995.

25. J. R. Tollervey, T. Curk, B. Rogelj, M. Briese, M. Cereda, M. Kayikci, J. König, T. Hortobágyi, A. L. Nishimura, V. Zupunski, R. Patani, S. Chandran, G. Rot, B. Zupan, C. E. Shaw, J. Ule, Characterizing the RNA targets and position-dependent splicing regulation by TDP-43. Nat. Neurosci. 14, 452–458 (2011). doi:10.1038/nn.2778.

26. M. J. Winton, L. M. Igaz, M. M. Wong, L. K. Kwong, J. Q. Trojanowski, V. M.-Y. Lee, Disturbance of nuclear and cytoplasmic TAR DNA-binding protein (TDP-43) induces disease-like redistribution, sequestration, and aggregate formation. J. Biol. Chem. 283, 13302–13309 (2008). doi:10.1074/jbc.M800342200.

27. M. Neumann, D. M. Sampathu, L. K. Kwong, A. C. Truax, M. C. Micsenyi, T. T. Chou, J. Bruce, T. Schuck, M. Grossman, C. M. Clark, L. F. McCluskey, B. L. Miller, E. Masliah, I. R. Mackenzie, H. Feldman, W. Feiden, H. A. Kretzschmar, J. Q. Trojanowski, V. M.-Y. Lee, Ubiquitinated TDP-43 in frontotemporal lobar degeneration and amyotrophic lateral sclerosis. Science 314, 130–133 (2006). doi:10.1126/science.1134108.

28. M. Polymenidou, C. Lagier-Tourenne, K. R. Hutt, S. C. Huelga, J. Moran, T. Y. Liang, S.-C. Ling, E. Sun, E. Wancewicz, C. Mazur, H. Kordasiewicz, Y. Sedaghat, J. P. Donohue, L. Shiue, C. F. Bennett, G. W. Yeo, D. W. Cleveland, Long pre-mRNA depletion and RNA missplicing contribute to neuronal vulnerability from loss of TDP-43. Nat. Neurosci. 14, 459–468 (2011). doi:10.1038/nn.2779.

29. T. Afroz, E.-M. Hock, P. Ernst, C. Foglieni, M. Jambeau, L. A. B. Gilhespy, F. Laferriere, Z. Maniecka, A. Plückthun, P. Mittl, P. Paganetti, F. H.-T. Allain, M. Polymenidou, Functional and dynamic polymerization of the ALS-linked protein TDP-43 antagonizes its pathologic aggregation. Nat. Commun. 8, 45 (2017). doi:10.1038/s41467-017-00062-0.

30. L. L. Jiang, W. Xue, J. Y. Hong, J. T. Zhang, M. J. Li, S. N. Yu, J. H. He, H. Y. Hu, The N-terminal dimerization is required for TDP-43 splicing activity. Sci. Rep. 7, 6196 (2017). doi:10.1038/s41598-017-06263-3.

31. M. Pérez-Berlanga, V. I. Wiersma, A. Zbinden, L. De Vos, U. Wagner, C. Foglieni, I. Mallona, K. M. Betz, A. Cléry, J. Weber, Z. Guo, R. Rigort, P. de Rossi, R. Manglunia, E. Tantardini, S. Sahadevan, O. Stach, M. Hruska-Plochan, F. H.-T. Allain, P. Paganetti, M. Polymenidou, Loss of TDP-43 oligomerization or RNA binding elicits distinct aggregation patterns. EMBO J. 42, e111719 (2023). doi:10.15252/embj.2022111719.

32. K. Oiwa, S. Watanabe, K. Onodera, Y. Iguchi, Y. Kinoshita, O. Komine, A. Sobue, Y. Okada, M. Katsuno, K. Yamanaka, Monomerization of TDP-43 is a key determinant for inducing TDP-43 pathology in amyotrophic lateral sclerosis. Sci. Adv. 9, eadf6895 (2023). doi:10.1126/sciadv.adf6895.

33. A. Wang, A. E. Conicella, H. B. Schmidt, E. W. Martin, S. N. Rhoads, A. N. Reeb, A. Nourse, D. Ramirez Montero, V. H. Ryan, R. Rohatgi, F. Shewmaker, M. T. Naik, T. Mittag, Y. M. Ayala, N. L. Fawzi, A single N-terminal phosphomimic disrupts TDP-43 polymerization, phase separation, and RNA splicing. EMBO J. 37, e97452 (2018). doi:10.15252/embj.201797452.

34. Ü. A. Laskay, A. A. Lobas, K. Srzentić, M. V. Gorshkov, Y. O. Tsybin, Proteome digestion specificity analysis for rational design of extended bottom-up and middle-down proteomics experiments. J. Proteome Res. 12, 5558–5569 (2013). doi:10.1021/pr400522h.

35. A. Leitner, R. Reischl, T. Walzthoeni, F. Herzog, S. Bohn, F. Förster, R. Aebersold, Expanding the chemical cross-linking toolbox by the use of multiple proteases and enrichment by size exclusion chromatography. Mol. Cell. Proteomics 11, M111.014126 (2012). doi:10.1074/mcp.M111.014126.

36. T. Dau, K. Gupta, I. Berger, J. Rappsilber, Sequential digestion with trypsin and elastase in cross-linking mass spectrometry. Anal. Chem. 91, 4472–4478 (2019). doi:10.1021/acs.analchem.8b05222.

37. Z. Ser, P. Cifani, A. Kentsis, Optimized cross-linking mass spectrometry for in situ interaction proteomics. J. Proteome Res. 18, 2545–2558 (2019). doi:10.1021/acs.jproteome.9b00085.

38. K. R. Adoni, J. E. Ditcham, A. K. González Rivera, G. H. Charlton, S. V. Saveliev, K. Thalassinos, R. Zenezini Chiozzi, Enhanced proteomics analysis with a novel recombinant chymotrypsin analogue engineered for high cleavage specificity. J. Proteome Res. 25, 2511–2519 (2026). doi:10.1021/acs.jproteome.5c01262.

39. S. A. McLuckey, G. J. Van Berkel, G. L. Glish, Tandem mass spectrometry of small, multiply charged oligonucleotides. J. Am. Soc. Mass Spectrom. 3, 60–70 (1992). doi:10.1016/1044-0305(92)85019-G.

40. M. G. Herrera, M. J. Amundarain, J. Santos, Biophysical evaluation of the oligomerization and conformational properties of the N-terminal domain of TDP-43. Arch. Biochem. Biophys. 737, 109533 (2023). doi:10.1016/j.abb.2023.109533.

41. M. Mompeán, V. Romano, D. Pantoja-Uceda, C. Stuani, F. E. Baralle, E. Buratti, D. V. Laurents, Point mutations in the N-terminal domain of transactive response DNA-binding protein 43 kDa (TDP-43) compromise its stability, dimerization, and functions. J. Biol. Chem. 292, 11992–12006 (2017). doi:10.1074/jbc.M117.775965.

42. T. Battellino, D. Yeung, H. Neustaeter, V. Spicer, K. Ogata, Y. Ishihama, O. V. Krokhin, Retention time prediction for post-translationally modified peptides: Ser, Thr, Tyr-phosphorylation. J. Chromatogr. A 1718, 464714 (2024). doi:10.1016/j.chroma.2024.464714.

43. J. Kim, K. Petritis, Y. Shen, D. G. Camp II, R. J. Moore, R. D. Smith, Phosphopeptide elution times in reversed-phase liquid chromatography. J. Chromatogr. A 1172, 9–18 (2007). doi:10.1016/j.chroma.2007.09.032.

44. D. Lal, E. M. Reinthaler, J. Altmüller, M. R. Toliat, H. Thiele, P. Nürnberg, H. Lerche, A. Hahn, R. S. Møller, H. Muhle, T. Sander, F. Zimprich, B. A. Neubauer, RBFOX1 and RBFOX3 mutations in Rolandic epilepsy. PLoS ONE 8, e73323 (2013). doi:10.1371/journal.pone.0073323.

45. J.-A. Lee, A. Damianov, C.-H. Lin, M. Fontes, N. N. Parikshak, E. S. Anderson, D. H. Geschwind, D. L. Black, K. C. Martin, Cytoplasmic Rbfox1 regulates the expression of synaptic and autism-related genes. Neuron 89, 113–128 (2016). doi:10.1016/j.neuron.2015.11.025.

46. G. Gómez, M. J. Pikal, N. Rodríguez-Hornedo, Effect of initial buffer composition on pH changes during far-from-equilibrium freezing of sodium phosphate buffer solutions. Pharm. Res. 18, 90–97 (2001). doi:10.1023/A:1011082911917.

47. C. Lenz, E. Kühn-Hölsken, H. Urlaub, Detection of protein-RNA crosslinks by nanoLC-ESI-MS/MS using precursor ion scanning and multiple reaction monitoring (MRM) experiments. J. Am. Soc. Mass Spectrom. 18, 869–881 (2007). doi:10.1016/j.jasms.2007.01.013.

48. W. Wang, L. Wang, J. Lu, S. L. Siedlak, H. Fujioka, J. Liang, S. Jiang, X. Ma, Z. Jiang, E. Lummertz da Rocha, M. Sheng, H. Choi, P. H. Lerou, H. Li, X. Wang, The inhibition of TDP-43 mitochondrial localization blocks its neuronal toxicity. Nat. Med. 22, 869–878 (2016). doi:10.1038/nm.4130.

49. L. François-Moutal, S. Perez-Miller, D. D. Scott, V. G. Miranda, N. Mollasalehi, M. Khanna, Structural insights into TDP-43 and effects of post-translational modifications. Front. Mol. Neurosci. 12, 301 (2019). doi:10.3389/fnmol.2019.00301.

50. K. C. Smith, R. T. Aplin, A mixed photoproduct of uracil and cysteine (5-S-cysteine-6-hydrouracil). A possible model for the in vivo cross-linking of deoxyribonucleic acid and protein by ultraviolet light. Biochemistry 5, 2125–2130 (1966). doi:10.1021/bi00870a046.

51. T. Jellinek, R. B. Johns, The mechanism of photochemical addition of cysteine to uracil and formation of dihydrouracil. Photochem. Photobiol. 11, 349–359 (1970). doi:10.1111/j.1751-1097.1970.tb06007.x.

52. C. L. Hawkins, M. J. Davies, Generation and propagation of radical reactions on proteins. Biochim. Biophys. Acta Bioenerg. 1504, 196–219 (2001). doi:10.1016/S0005-2728(00)00252-8.

53. H. Feng, X.-J. Lu, S. Maji, L. Liu, D. Ustianenko, N. D. Rudnick, C. Zhang, Structure-based prediction and characterization of photo-crosslinking in native protein–RNA complexes. Nat. Commun. 15, 2279 (2024). doi:10.1038/s41467-024-46429-y.

54. B. J. Calnan, B. Tidor, S. Biancalana, D. Hudson, A. D. Frankel, Arginine-mediated RNA recognition: The arginine fork. Science 252, 1167–1171 (1991). doi:10.1126/science.252.5009.1167.

55. S. S. Chavali, C. E. Cavender, D. H. Mathews, J. E. Wedekind, Arginine forks are a widespread motif to recognize phosphate backbones and guanine nucleobases in the RNA major groove. J. Am. Chem. Soc. 142, 19835–19839 (2020). doi:10.1021/jacs.0c09689.

56. Z. Yang, X. Deng, Y. Liu, W. Gong, C. Li, Analyses on clustering of the conserved residues at protein-RNA interfaces and its application in binding site identification. BMC Bioinformatics 21, 57 (2020). doi:10.1186/s12859-020-3398-9.

57. R. L. French, Z. R. Grese, H. Aligireddy, D. D. Dhavale, A. N. Reeb, N. Kedia, P. T. Kotzbauer, J. Bieschke, Y. M. Ayala, Detection of TAR DNA-binding protein 43 (TDP-43) oligomers as initial intermediate species during aggregate formation. J. Biol. Chem. 294, 6696–6709 (2019). doi:10.1074/jbc.RA118.005889.

58. J. R. Mann, A. M. Gleixner, J. C. Mauna, E. Gomes, M. R. DeChellis-Marks, P. G. Needham, K. E. Copley, B. Hurtle, B. Portz, N. J. Pyles, L. Guo, C. B. Calder, Z. P. Wills, U. B. Pandey, J. K. Kofler, J. L. Brodsky, A. Thathiah, J. Shorter, C. J. Donnelly, RNA binding antagonizes neurotoxic phase transitions of TDP-43. Neuron 102, 321–338 (2019). doi:10.1016/j.neuron.2019.01.048.

59. W. H. Hudson, E. A. Ortlund, The structure, function and evolution of proteins that bind DNA and RNA. Nat. Rev. Mol. Cell Biol. 15, 749–760 (2014). doi:10.1038/nrm3884.

60. M. D. Shetlar, Cross-linking of proteins to nucleic acids by ultraviolet light, in Photochemical and Photobiological Reviews, K. C. Smith, Ed., Springer (1980), vol. 5, pp. 105–197. doi:10.1007/978-1-4684-3641-9_4.

61. T. R. Stachowski, M. Fischer, Unfreezing structural biology for drug discovery. Nat. Chem. Biol. 22, 169–179 (2026). doi:10.1038/s41589-025-02047-3.

62. C. H. Vieira-Vieira, M. Selbach, Opportunities and challenges in global quantification of RNA-protein interaction via UV cross-linking. Front. Mol. Biosci. 8, 669939 (2021). doi:10.3389/fmolb.2021.669939.

63. A. Panhale, F. M. Richter, F. Ramírez, M. Shvedunova, T. Manke, G. Mittler, A. Akhtar, CAPRI enables comparison of evolutionarily conserved RNA interacting regions. Nat. Commun. 10, 2682 (2019). doi:10.1038/s41467-019-10585-3.

64. C. P. Sarnowski, M. Götze, A. Leitner, RNxQuest: An extension to the xQuest pipeline enabling analysis of protein–RNA cross-linking/mass spectrometry data. J. Proteome Res. 22, 3368–3382 (2023). doi:10.1021/acs.jproteome.3c00341.

65. J. W. Bae, S. C. Kwon, Y. Na, V. N. Kim, J.-S. Kim, Chemical RNA digestion enables robust RNA-binding site mapping at single amino acid resolution. Nat. Struct. Mol. Biol. 27, 678–682 (2020). doi:10.1038/s41594-020-0436-2.

66. B. Bogdanow, H. Zauber, M. Selbach, Systematic errors in peptide and protein identification and quantification by modified peptides. Mol. Cell. Proteomics 15, 2791–2801 (2016). doi:10.1074/mcp.M115.055103.

67. E. Gasteiger, C. Hoogland, A. Gattiker, S. Duvaud, M. R. Wilkins, R. D. Appel, A. Bairoch, Protein identification and analysis tools on the ExPASy server, in The Proteomics Protocols Handbook, J. M. Walker, Ed., Humana Press (2005), pp. 571–607. doi:10.1385/1592598900.

68. M. C. Chambers, B. MacLean, R. Burke, D. Amodei, D. L. Ruderman, S. Neumann, L. Gatto, B. Fischer, B. Pratt, J. Egertson, K. Hoff, D. Kessner, N. Tasman, N. Shulman, B. Frewen, T. A. Baker, M.-Y. Brusniak, C. Paulse, D. Creasy, L. Flashner, K. Kani, C. Moulding, S. L. Seymour, L. M. Nuwaysir, B. Lefebvre, F. Kuhlmann, J. Roark, R. Paape, D. Suckau, T. Hemenway, A. Huhmer, J. Langridge, B. Connolly, T. Chadick, K. Holly, J. Eckels, E. W. Deutsch, R. L. Moritz, J. E. Katz, D. B. Agus, M. MacCoss, D. L. Tabb, P. Mallick, A cross-platform toolkit for mass spectrometry and proteomics. Nat. Biotechnol. 30, 918–920 (2012). doi:10.1038/nbt.2377.

69. O. Rinner, J. Seebacher, T. Walzthoeni, L. N. Mueller, M. Beck, A. Schmidt, M. Mueller, R. Aebersold, Identification of cross-linked peptides from large sequence databases. Nat. Methods 5, 315–318 (2008). doi:10.1038/nmeth.1192.

70. T. Walzthoeni, M. Claassen, A. Leitner, F. Herzog, S. Bohn, F. Förster, M. Beck, R. Aebersold, False discovery rate estimation for cross-linked peptides identified by mass spectrometry. Nat. Methods 9, 901–903 (2012). doi:10.1038/nmeth.2103.

71. B. MacLean, D. M. Tomazela, N. Shulman, M. Chambers, G. L. Finney, B. Frewen, R. Kern, D. L. Tabb, D. C. Liebler, M. J. MacCoss, Skyline: An open source document editor for creating and analyzing targeted proteomics experiments. Bioinformatics 26, 966–968 (2010). doi:10.1093/bioinformatics/btq054.

72. C. Fu, X. Zheng, Y. Jiang, Y. Liu, P. Xu, Z. Zeng, R. Liu, Y. Zhao, A universal and multiplex kinase assay using γ-[18O4]-ATP. Chem. Commun. 49, 2795–2797 (2013). doi:10.1039/c3cc38467e.

73. J. Alguacil, D. Reyes, Y. Aubin, B. Roy, C. Périgaud, E. Champagne, S. Peyrottes, Exploring synthetic pathways for nucleosidic derivatives of potent phosphoantigens. New J. Chem. 40, 6046– 6055 (2016). doi:10.1039/c5nj03614c.

