## Supplementary information for "A qualitative and quantitative framework for the characterization of protein–RNA interactions by cross-linking mass spectrometry (XL-MS)"

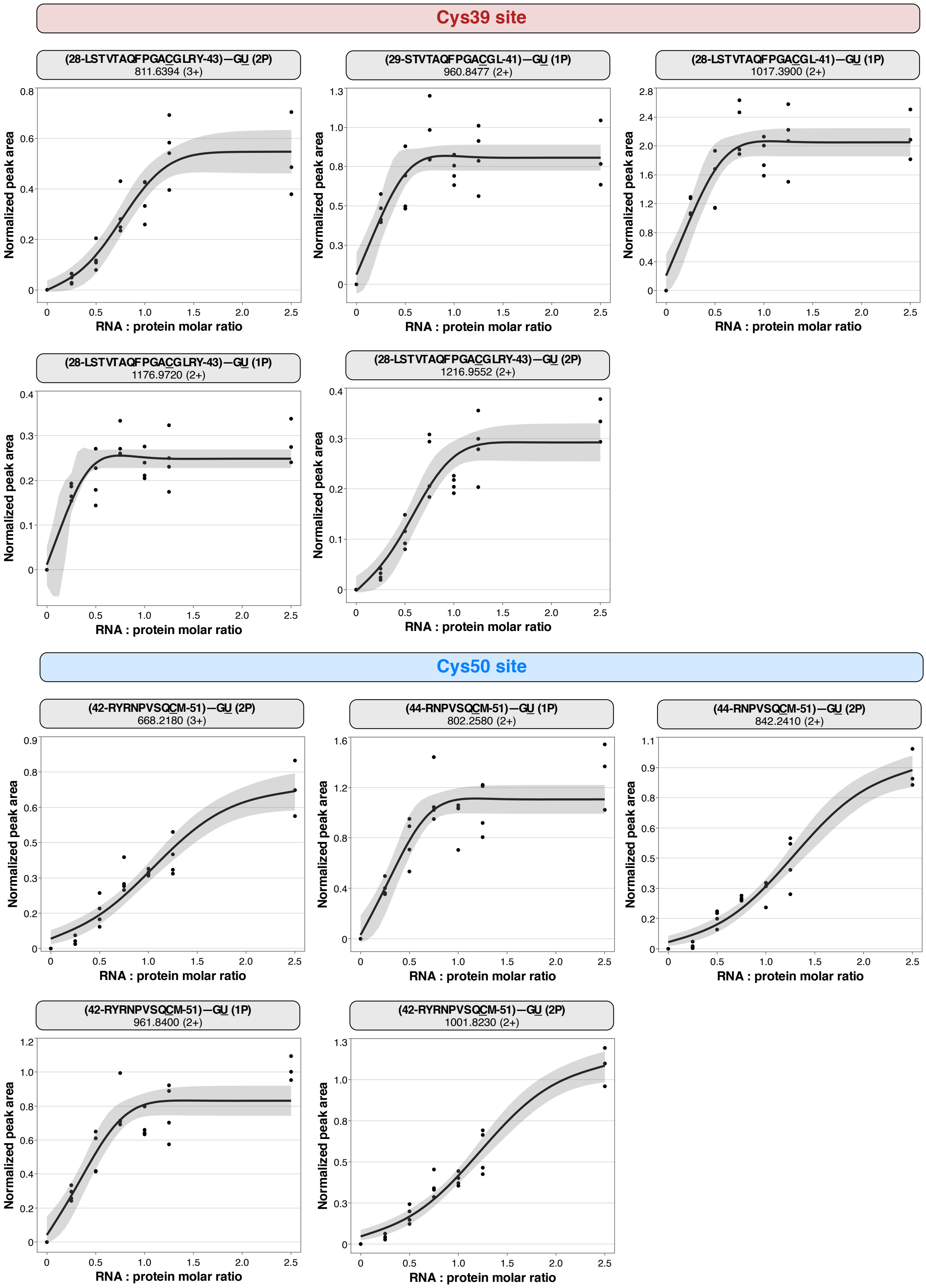

**Supplementary Figure 1.** Individual RNA titration curves obtained for the cross-links reporting on the Cys39 (left) and on the Cys50 interaction site (right) in TDP-43 NTD. Peptide and RNA residues involved in cross-link formation are underlined. Normalized peak area data points across RNA–protein molar ratios were fitted to a three-parameter equation. The 95% confidence band around each fit represents mean predicted response. 1P: phosphate-lacking form, 2P: phosphate-retaining form of a cross-link.

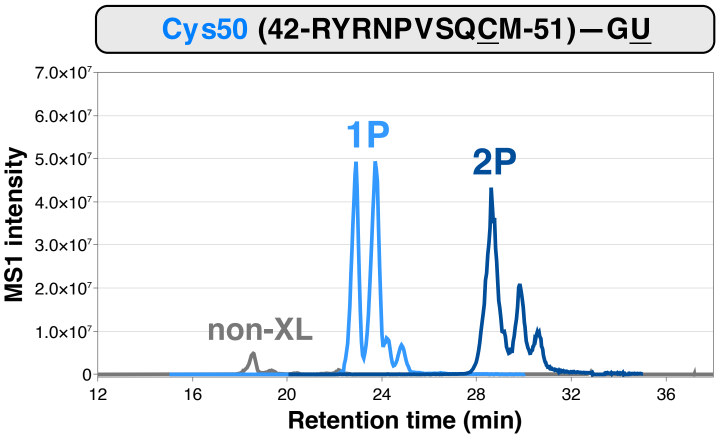


**Supplementary Figure 2.** Retention time comparison of the RYRNPVSQCM–GU peptide–RNA cross-link of TDP-43 NTD WT. Peptide and RNA residues involved in cross-link formation are underlined. GU denotes the RNA composition, rather than the sequence. Shown are the non-cross-linked peptide (grey) and the cross-linked forms of the same peptide with RNA lacking one phosphate group (1P, light blue) or retaining it (2P, dark blue). Data shown on the example of one of the replicates of RNA-to-protein ratio of 1. The observed retention pattern is consistent with the other measured RNA-to-protein ratios.

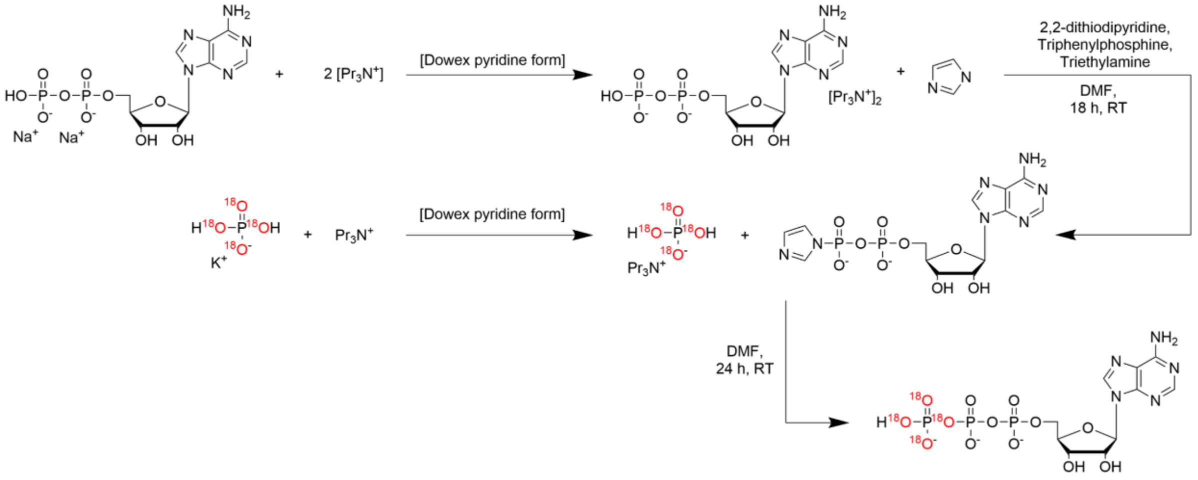


**Supplementary Figure 3.** A schematic overview of the $^{18}O_4$-γ-ATP synthesis.

| TDP-43 NTD variants comparison experiment | | | | | | |
|---|---|---|---|---|---|---|
| **Cross-link** | **m/z light state** | **m/z heavy state** | **Charge state** | **Retention time window (min)** | **Phosphorylation form** | **Neutral loss** |
| RNPVSQCM–GU | 535.1740 | 537.1790 | 3+ | 15–35 | 1P | -$HPO_3$ |
| LSTVTAQFPGACGL–GU | 678.5960 | 680.6000 | 3+ | 45–65 | 1P | -$HPO_3$ |
| RNPVSQCM–GU | 802.2580 | 805.2640 | 2+ | 15–35 | 1P | -$HPO_3$ |
| STVTAQFPGACGL–GU | 960.8480 | 963.8540 | 2+ | 45–65 | 1P | -$HPO_3$ |
| RYRNPVSQCM–GU | 1001.8230 | 1004.8290 | 2+ | 15–35 | 2P | |
| LSTVTAQFPGACGL–GU | 1017.3900 | 1020.3960 | 2+ | 45–65 | 1P | -$HPO_3$ |
| RYRNPVSQCM–GU | 668.2180 | 670.2220 | 3+ | 15–35 | 2P | |
| RYRNPVSQCM–GU | 641.5620 | 643.5670 | 3+ | 15–35 | 1P | -$HPO_3$ |
| RYRNPVSQCM–GU | 961.8400 | 964.8460 | 2+ | 15–35 | 1P | -$HPO_3$ |
| RNPVSQCM–GU | 842.2410 | 845.2470 | 2+ | 15–35 | 2P | -$HPO_3$ |

| TDP-43 NTD WT–$(GU)_6$ titration experiment | | | | | | |
|---|---|---|---|---|---|---|
| **Cross-link** | **m/z light state** | **m/z heavy state** | **Charge state** | **Retention time window (min)** | **Phosphorylation form** | **Neutral loss** |
| RYRNPVSQCM–GU | 641.5624 | 643.5667 | 3+ | 15–30 | 1P | -$HPO_3$ |
| RYRNPVSQCM–GU | 668.2179 | 670.2221 | 3+ | 22–37 | 2P | |
| RNPVSQCM–GU | 802.2575 | 805.2639 | 2+ | 15–30 | 1P | -$HPO_3$ |
| LSTVTAQFPGACGLRY–GU | 811.6394 | 813.6436 | 3+ | 50–65 | 2P | |
| RNPVSQCM–GU | 842.2407 | 845.2470 | 2+ | 20–35 | 2P | |
| STVTAQFPGACGL | 960.8477 | 963.8541 | 2+ | 45–60 | 1P | -$HPO_3$ |
| RYRNPVSQCM–GU | 961.8397 | 964.8461 | 2+ | 15–30 | 1P | -$HPO_3$ |
| RYRNPVSQCM–GU | 1001.8229 | 1004.8293 | 2+ | 20–35 | 2P | |
| LSTVTAQFPGACGL–GU | 1017.3898 | 1020.3961 | 2+ | 45–65 | 1P | -$HPO_3$ |
| LSTVTAQFPGACGLRY–GU | 1176.9720 | 1179.9784 | 2+ | 40–55 | 1P | -$HPO_3$ |
| LSTVTAQFPGACGLRY–GU | 1216.9552 | 1219.9615 | 2+ | 50–65 | 2P | -$HPO_3$ |

| FOX1-RRM experiment | | | | | | |
|---|---|---|---|---|---|---|
| **Cross-link** | **m/z light state** | **m/z heavy state** | **Charge state** | **Retention time window (min)** | **Phosphorylation form** | **Neutral loss** |
| LHVSNIPFR–CG | 583.5700 | 585.574 | 3+ | 10–18 | 1P | $-H_3PO_3$ |
| RLHVSNIPFR–AU | 657.9280 | 659.932 | 3+ | 24–32 | 2P | |
| LHVSNIPFR–CGU | 685.5790 | 687.583 | 3+ | 12–30 | 1P | $-H_3PO_3$ |
| GFGFVTFENSADADR–G | 691.9180 | 693.922 | 3+ | 10–27 | 2P | $-H_2$ |
| GFGFVTFENSADADR–G | 692.5900 | 694.594 | 3+ | 10–27 | 2P | |
| LHVSNIPFR–CGU | 712.2340 | 714.238 | 3+ | 15–25 | 2P | $-H_2$ |
| LHVSNIPFR–CGU | 712.9060 | 714.910 | 3+ | 15–25 | 2P | |
| GFGFVTFENSADADR–GU | 773.9460 | 775.951 | 3+ | 26–33 | 1P | $-HPO_3+H_2O$ |
| LHVSNIPFRFR–CGU | 813.9620 | 815.967 | 3+ | 20–28 | 2P | |
| GFGFVTFENSADADR–CGU | 875.6270 | 877.631 | 3+ | 26–34 | 1P | $-HPO_3+H_2O$ |
| GFGFVTFENSADADR–AGU | 877.6270 | 879.631 | 3+ | 28–36 | 1P | $-HPO_3$ |
| GFGFVTFENSADADR–AGU | 883.6305 | 885.635 | 3+ | 25–34 | 1P | $-HPO_3+H_2O$ |
| GFGFVTFENSADADR–AGU | 903.6105 | 905.615 | 3+ | 36–42 | 2P | $-H_2$ |
| GFGFVTFENSADADR–AGU | 904.2824 | 906.287 | 3+ | 36–42 | 2P | |
| LHVSNIPFR–GU | 915.3264 | 918.333 | 2+ | 12–30 | 2P | $-H_2$ |
| LHVSNIPFR–GU | 916.3343 | 919.341 | 2+ | 12–30 | 2P | |
| GFGFVTFENSADADR–AGUU | 978.9635 | 980.968 | 3+ | 35–48 | 1P | $-H_3PO_3$ |
| GFGFVTFENSADADR–AGUU | 979.6354 | 981.640 | 3+ | 35–48 | 1P | $-HPO_3$ |
| GFGFVTFENSADADR–AGUU | 985.6389 | 987.643 | 3+ | 30–48 | 1P | $-HPO_3+H_2O$ |
| GFGFVTFENSADADR–GU | 1150.4025 | 1153.409 | 2+ | 28–36 | 1P | $-H_3PO_3$ |
| GFGFVTFENSADADR–GU | 1159.4077 | 1162.414 | 2+ | 28–36 | 1P | $-H_3PO_3+H_2O$ |
| GFGFVTFENSADADR–AGU | 1314.9287 | 1317.935 | 2+ | 28–36 | 1P | $-H_3PO_3$ |

**Supplementary Table 1.** Inclusion lists of peptide-RNA cross-links targeted in the PRM experiments performed in this study. Wide retention time windows were used due to broad elution profiles of the target cross-links and their appearance as several chromatographic peaks. 1P: phosphate-lacking form, 2P: phosphate-retaining form of a cross-link.
